# The publication and citation gender gap in Biology: bibliometric analysis of Biology faculty at top United States universities

**DOI:** 10.1101/2025.09.30.679585

**Authors:** David Alvarez-Ponce, James Vesper

## Abstract

Women are underrepresented in academia, especially in top research institutions, STEM departments, and senior positions. A number of studies have shown that, on average, female academics publish fewer articles per year, are less cited, and are promoted more slowly than their male peers. Other studies, however, have found that these gender differences are absent or even reversed in certain fields and social contexts, highlighting the importance of evaluating the different dimensions of the gender gap separately within each field and context. We obtained a census of 2104 female and 3721 male tenured and tenure-eligible faculty members affiliated with Biology Departments at the 146 United States R1 universities. We show that women represent 46.94% of Assistant Professors, 38.51% of Associate Professors and 30.09% of Professors in our dataset. Compared with their male peers, female faculty members tend to produce fewer publications per year (even after controlling for career stage and university ranking), to publish in lower-impact factor journals, and to be less cited (even after controlling for career length, career stage, number of publications, average impact factor of the journals in which they publish, and university ranking). Despite their lower publication and citation rates, female faculty members in our dataset require the same amount of time to attain the rank of Professor as their male peers.

## INTRODUCTION

Women are underrepresented in academia, especially in top research institutions, STEM departments, and senior positions (Baker and Koedel 2024; Ceci et al. 2009, 2014; Ginther and Kahn 2004; Holman et al. 2018; Hornig 1980; Rushworth et al. 2021). This underrepresentation can be attributed to a combination of factors. First, even though the number of women entering academia has considerably increased over the last decades, it was very low in the past, resulting in few women currently occupying senior positions (Alvarez-Ponce and Vesper 2025; Ceci et al. 2014; Fox et al. 2019; Holman et al. 2018). Second, women are more likely than men to leave academia, either voluntarily or involuntarily, resulting in shorter academic careers and contributing to women’s underrepresentation at senior positions (Goulden et al. 2011; Shaw and Stanton 2012). Third, women seem to obtain tenure and to be promoted more slowly than their male peers (Ginther and Kahn 2004; Heijstra et al. 2015; Long et al. 1993; Weisshaar 2017).

In many academic fields there is a gender productivity gap: on average, female researchers produce fewer publications, which hinders their career advancement (Astegiano et al. 2019; J. R. Cole 2024; Holman et al. 2018; Larivière et al. 2013; Martin 2012; West et al. 2013). In addition, in many academic fields, women receive less recognition from their peers, e.g., in the form of authorship (Ross et al. 2022), research funding (Goulden et al. 2011; Martin 2012), invitations to present their work at conferences and publish their research in journals (Holman et al. 2018; Schroeder et al. 2013), and citations (Chatterjee and Werner 2021; Davenport and Snyder 1995; Ferber and Brün 2011; Larivière et al. 2013; Maliniak et al. 2013; Teich et al. 2022), which further limits their opportunities for career advancement.

Even though the productivity gap seems to be pervasive, its severity depends on the specific social context (including the specific discipline, country, kind of institution, and job title), being absent or even reversed (i.e., women publish at higher rates) in certain settings (Arensbergen et al. 2012; Duch et al. 2012; Frandsen et al. 2020; Schucan Bird 2011). Some analyses have attributed the productivity gap almost entirely to the fact that women tend to have shorter careers (Huang et al. 2020), whereas others have shown that female academics tend to publish less than their male peers even after controlling for the length of their careers (i.e., they produce on average fewer publications per year; e.g. Duch et al. 2012).

The reasons for the productivity gap remain unclear, which has led to researchers coining the term “productivity puzzle” (J. R. Cole 2024; J. Cole and Zuckerman 1984), but some explanations have been proposed. First, female academics tend to secure less funding than their male peers, which limits their ability to produce research (Goulden et al. 2011; Martin 2012); in fact, the gender productivity gap is smaller in fields that require less funding (Duch et al. 2012). Second, female academics often assume a greater share of childcare and home responsibilities, which limits the amount of time that they can devote to their careers (Awung and Dorasamy 2015; Ledin et al. 2007; Rhoads and Rhoads 2012). Third, female academics tend to have higher teaching and/or service loads than their male peers, which limits the amount of effort that they can devote to their research (Eagly 2020; Guarino and Borden 2017; O’Meara et al. 2017). Fourth, women might be more prone to perfectionism, risk aversion, self-doubt and self-criticism (Born et al. 2022; Coffman 2014; Croson and Gneezy 2009; Exley and Kessler 2022; Kessler et al. 2014; Möbius et al. 2022; Shastry and Shurchkov 2024), which results in them submitting fewer manuscripts per year. Last, female-authored manuscripts may be held to higher standards by editors and/or referees (Card et al. 2020; Hengel 2022).

In addition to being evaluated based on their publication rates, researchers are often evaluated based on the impact of their publications. Commonly used, simple metrics of research impact include the total number of citations received by their publications and the average number of citations per publication. A more sophisticated metric is a researcher’s *h*-index, which is the maximum quantity (*h*) such that the researcher has published *h* publications with at least *h* citations each (Hirsch 2005). For instance, a researcher with 30 publications, including 15 publications with 15 or more citations each, will have an *h*-index of 15. An advantage of the *h*-index is that it is not significantly influenced by outliers (very highly cited publications). However, it strongly correlates with a researcher’s career length (i.e., with the number of years since their first publication), which led to formulating another metric: the *m*-index, which is a researcher’s *h*-index divided by the number of years since their first publication (Hirsch 2005). For instance, a researcher with an *h*-index of 15 that has been publishing for 10 years will have an *m*-index of 15/10 = 1.5.

Some analyses have suggested that male researchers tend to be more cited (Chatterjee and Werner 2021; Davenport and Snyder 1995; Ferber and Brün 2011; Larivière et al. 2013; Maliniak et al. 2013; Teich et al. 2022), which could be due to: (1) their higher productivity (researchers tend to cite scientists that they encounter more often in the literature—the so-called “fast-food effect”; Kelly and Jennions 2006); (2) them enjoying, on average, a higher standing in the academic community (contributions introduced by prestigious academics tend to enjoy more visibility—the so-called “Mathew effect”; Merton 1968); (3) them more often publishing in highly prestigious journals (Bendels et al. 2018; Krishnamurthy et al. 2017; Y. A. Shen et al. 2018), (4) them self-citing more often, which has both direct and indirect effects on their citation metrics (Fowler and Aksnes 2007); (5) them having larger co-author networks (Bosquet and Combes 2013; Ductor et al. 2023; Mcdowell and Smith 1992; Rothstein and Davey 1995); or (6) a “lottery effect”—by publishing more articles, male authors are more likely to publish highly-cited articles by chance (Kelly and Jennions 2006). However, other analyses have obtained contrasting results: female-authored articles tend to be more cited than male-authored ones in certain contexts (Card et al. 2020; Duch et al. 2012; Long 1992; Symonds et al. 2006), which could be due to them being of higher quality—perhaps as a result to them being held to higher standards during peer review (Card et al. 2020). Other analyses, on the other hand, have found no significant differences in the citations received by male-and female-authored research articles after removing the effect of male authors’ higher productivities (Aksnes et al. 2011; Huang et al. 2020; Pautasso 2013).

Typical publication and citation practices and metrics are specific to each field, country and kind of institution (e.g., Alonso et al. 2009; Hirsch 2005). In addition, they have significantly changed over time. This probably explains why findings from different studies can contradict each other, and highlights the importance of evaluating the gender gap within each field and social context separately. Here, we compare the research outputs of 2104 female and 3721 male tenure-eligible (Assistant Professors) and tenured (Associate Professors or Professors) faculty members working in the Biology departments of the 146 United States universities classified in the R1 category (“Doctoral Universities – Very high research activity”) by the Carnegie Classification of Institutions of Higher Education. To our knowledge, no study has analyzed the research output of faculty in the field of Biology as a whole. In addition, our selection of faculty members offers some advantages. First, biologists, especially those at top institutions, produce a large percent of research articles published annually, both in the United States and worldwide (Baas et al. 2020; White 2019). Second, hiring and career advancement practices are relatively uniform across R1 universities: typically, new faculty members entering the so-called “tenure track” are first hired as Assistant Professors (who are often untenured); after approximately 5 years, they apply for promotion to Associate Professors (who are often tenured); later in their careers they may apply for promotion to Professors (also referred to as Full Professors). Third, Assistant Professors, Associate Professors and Professors at R1 universities often have similar responsibilities, which include research, teaching and service (Price and Cotten 2006).

We show that: (1) Women account for 46.94% of Assistant Professors, 38.51% of Associate Professors and 30.09% of Professors at R1 Biology departments. (2) Female representation is only slightly lower (45.79% of Assistant Professors, 36.70% of Associate Professors and 28.52% of Professors) at the Biology departments of the 50 top United States Universities. (3) Male faculty members tend to produce a higher number of publications per year than their female peers, even after controlling for career stage and university ranking. (4) Male faculty members tend to publish in higher-impact factor journals than their female peers. (5) Male faculty members tend to be more cited than their female peers, even after controlling for career length, career stage, number of publications, average impact factor of the journals in which they publish, and university ranking. (6) Despite their lower publication and citation rates, female faculty members in our dataset require the same amount of time to attain the rank of Professor as their male peers.

## METHODS

### List of Biology faculty members affiliated with United States R1 universities

We obtained a list of United States universities classified in the R1 category (i.e., the most research-intensive category) by the Carnegie Classification of Institutions of Higher Education (*n* = 146). The order of the universities was randomized before data collection. For each university, we obtained a list of Biology faculty from the relevant department or program websites. Most of these universities had a Biology department or program (common names include “Biology”, “Integrative Biology”, “Biological Sciences”, “Biosciences” and “Life Sciences”). Other universities had separate departments focusing on Ecology, Evolution and Conservation Biology and on Cellular and Molecular Biology, in which case all relevant departments were included. Only tenure-track (Assistant Professors) or tenured (Associate Professors and Professors) faculty members were included in our census. Only-research, only-teaching, only-service, clinical, practice, emeritus, adjunct and visiting faculty members were excluded in order to obtain a research-active set of faculty members with comparable duties. Genders were inferred from the information available on the department’s websites (pronouns and pictures). The list of faculty members was obtained between June 18 and July 10, 2024.

### Bibliometric data

For each faculty member, we obtained their total number of publications, total number of citations, *h*-index (Hirsch 2005), year of first publication, and full list of publications from their Scopus profile (Baas et al. 2020). Profiles were initially located using each individual’s name and institution. If our initial search failed or produced multiple profiles, we tried to locate the individual by comparison with their faculty or personal website (looking at their publications and previous affiliations). Individuals with multiple profiles were excluded, unless one of the profiles included all the publications included in the alternative profiles. Bibliometric data were collected between July 10 and August 31, 2024.

From the collected data, we computed: (1) the number of years since first publication, by subtracting the year of first publication from 2024; (2) the number of publications per year, by dividing the total number of publications by the number of years since first publication; (3) the number of citations per year, by dividing the total number of citations by the number of years since first publication; (4) the *m*-index, by dividing the *h*-index by the total number of years since first publication; and (5) the number of citations per publication, by dividing the total number of citations by the total number of publications.

In addition, for each author we parsed their list of publications to compute: (1) the number of publications in 2023; (2) the average number of authors per publication; (3) the size of their co-author network (i.e., the total number of individuals with whom they have co-authored publications at some point); (4) the percent of their publications of which they are the first author; (5) the percent of their publications of which they are the last author; (6) the percent of their publications of which they are sole author; and (7) the average journal impact factor across all their publications.

### Additional information

For each university, we obtained its ranking from the *U.S. News & World Report* Best Colleges Ranking (https://www.usnews.com/best-colleges/rankings/national-universities; last accessed on September 1, 2024). For each journal represented in our dataset, we retrieved its impact factor from the 2023 *Journal Citation Reports* (Clarivate, Philadelphia, PA).

### Statistical analyses

All statistical analyses were conducted using R version 4.4.2. For ANCOVA analyses, we used the rstatix package.

## RESULTS

### Gender composition of tenured and tenure-eligible Biology faculty at R1 universities

We assembled a list of 5825 Biology faculty members affiliated with the 146 United States universities classified in the R1 category. Our list included 2104 women and 3721 men. In terms of academic rank, the list included 1338 Assistant Professors (628 women and 710 men), 1493 Associate Professors (575 women and 918 men) and 2994 Professors (901 women and 2093 men; Fig. 1). Thus, women represent 36.12% of our sample (46.94% of Assistant Professors, 38.51% of Associate Professors and 30.09% of Professors).

**Figure 1:**
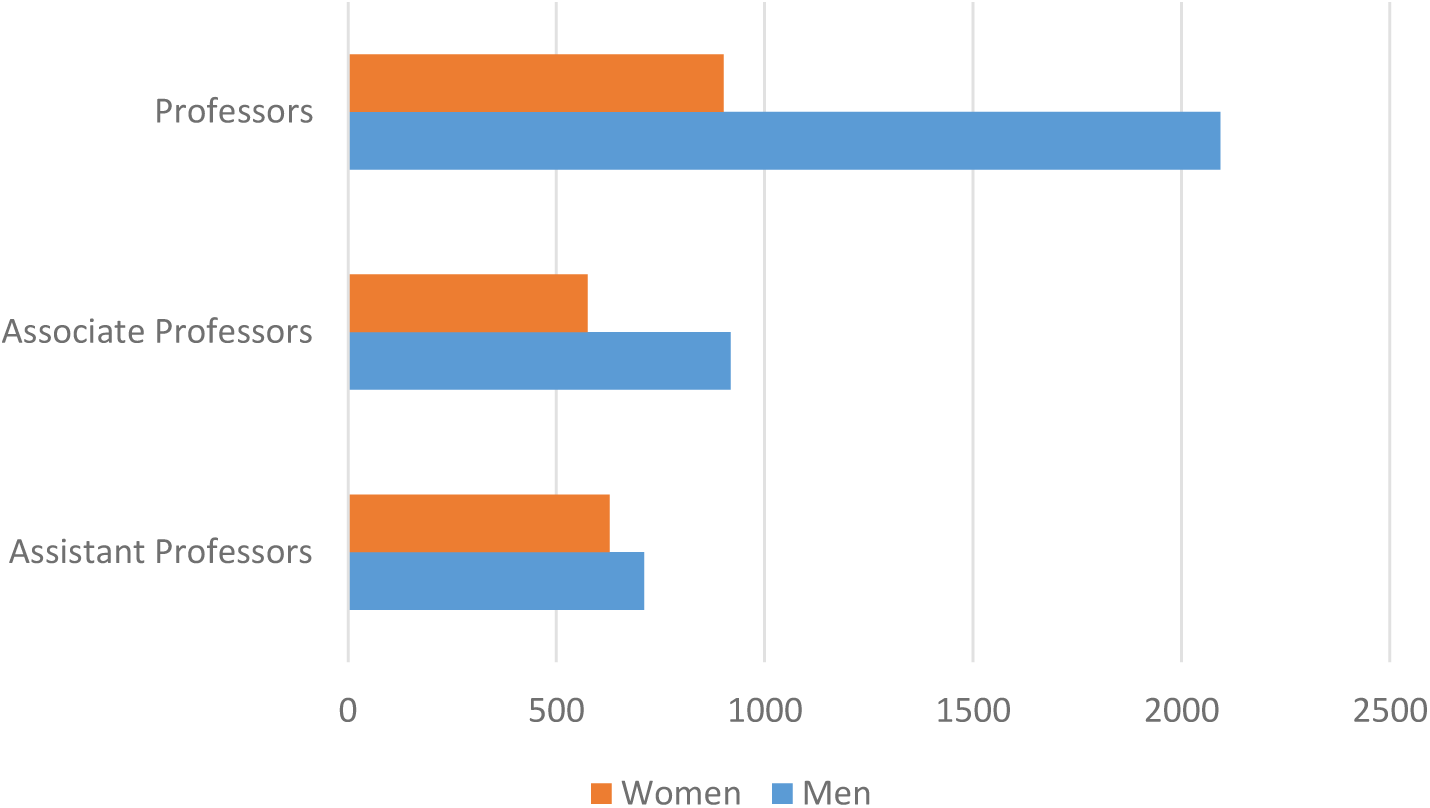
Gender and rank composition of Biology tenured and tenure-eligible faculty affiliated with United States R1 universities.

For each university, we computed the fraction of women among all faculty members (ranging from 18.18% to 71.43%), among Assistant Professors (0–100%), among Associate Professors (0– 75%) and among Professors (0–75%). We found a weak but significantly positive correlation between universities’ rankings and their fraction of female faculty (Spearman’s rank correlation coefficient; ρ = 0.172, *n* = 146, *P* = 0.038). However, the correlation was not significant for Assistant Professors (ρ = 0.055, *n* = 146, *P* = 0.509), Associate Professors (ρ = 0.037, *n* = 145, *P* = 0.663) or Professors (ρ = −0.024, *n* = 146, *P* = 0.778) separately. In addition, the faculty at the top 50 universities were 33.85% female (45.79% of Assistant Professors, 36.70% of Associate Professors and 28.52% of Professors). These results indicate that women are only slightly underrepresented in the Biology departments of the 50 top United States universities compared with the Biology departments of other R1 universities.

### Male faculty members publish at higher rates than female faculty members

We could find the Scopus profile for 5549 of the 5828 faculty members in our dataset (i.e., 95.21%). Table 1 shows the average and median total number of publications of faculty members in six different categories (female Assistant Professors, male Assistant Professors, female Associate Professors, male Associate Professors, female Professors, male Professors), and Fig. 2 represents the distribution of the number of publications for each category. As expected, Professors had on average more publications than Associate Professors (Mann–Whitney’s *U* test, *P* = 5.74×10^−266^) and Associate Professors had on average more publications than Assistant Professors (*P* = 1.15×10^−123^). Within each of the three academic ranks, men authored significantly more publications than women (Mann–Whitney’s *U* test, Assistant Professors: *P* = 9.46×10^−11^, Associate Professors: *P* = 1.17×10^−11^, Professors: *P* = 3.21×10^−20^).

**Figure 2:**
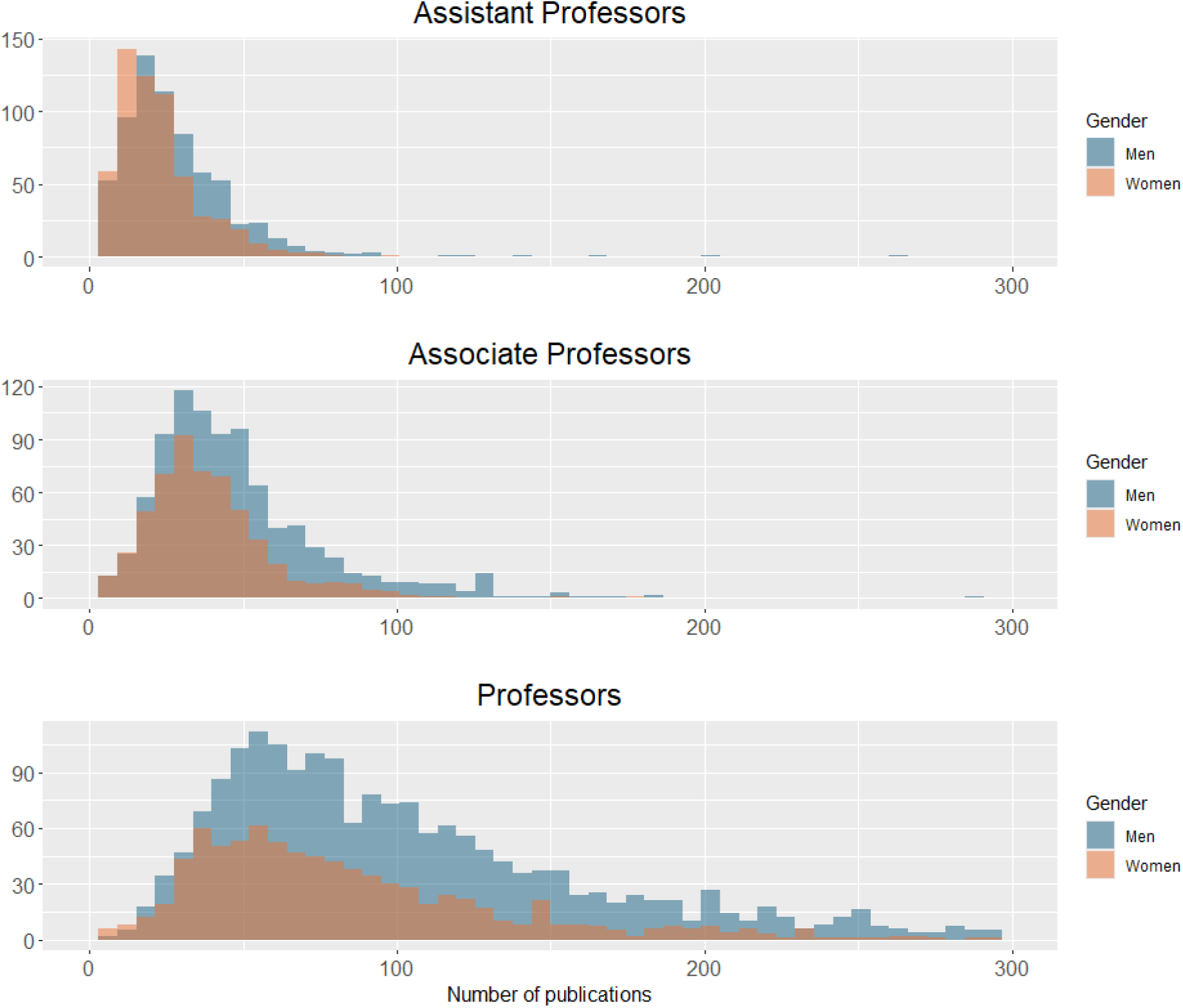
Number of publications of female and male faculty members of different ranks.

**Table 1:** Number of publications.

| Gender and rank | All 146 R1 universities |  |  | *P*-value<sup>a</sup> | Top 50 universities |  |  | *P*-value<sup>b</sup> | *P*-value<sup>c</sup> |
| --- | --- | --- | --- | --- | --- | --- | --- | --- | --- |
|  | *n* | Average | Median |  | *n* | Average | Median |  |  |
| Female Assistant Prof. | 628 | 22.07 | 19.0 | **9.46×10<sup>-11</sup>** | 250 | 23.29 | 20.0 | **0.0001** | 0.0748 |
| Male Assistant Prof. | 710 | 28.72 | 24.0 |  | 296 | 28.33 | 25.0 |  | 0.2848 |
| Female Associate Prof. | 575 | 37.96 | 35.0 | **1.17×10<sup>-11</sup>** | 178 | 44.90 | 41.0 | **0.0026** | **1.35×10<sup>-7</sup>** |
| Male Associate Prof. | 918 | 48.14 | 41.0 |  | 307 | 52.18 | 46.0 |  | **1.19×10<sup>-5</sup>** |
| Female Professors | 901 | 87.13 | 72.0 | **3.21×10<sup>-20</sup>** | 423 | 99.06 | 79.5 | **4.30×10<sup>-11</sup>** | **1.84×10<sup>-9</sup>** |
| Male Professors | 2093 | 119.10 | 91.0 |  | 1060 | 135.70 | 104.0 |  | **1.53×10<sup>-20</sup>** |

<sup>a</sup>Comparison of female vs. male faculty members affiliated with all 146 R1 universities.
<sup>b</sup>Comparison of female vs. male faculty members affiliated with the top 50 R1 universities.
<sup>c</sup>Comparison of faculty members affiliated with the top 50 R1 universities vs. faculty members affiliated with the other 96 R1 universities.
All *P*-values correspond to the Mann–Whitney *U* test. *P*-values lower than 0.05 are shown in bold face.

For each faculty member, we computed the time since their first publication by subtracting the year of their first publication from 2024. Table S1 shows the average and median values for faculty members in the six categories, and Fig. S1 shows the distribution for each category. As expected, on average Professors have been publishing for a longer amount of time than Associate Professors (Mann–Whitney’s *U* test, *P* = 8.43×10^−221^) and Associate Professors have been publishing for a longer amount of time than Assistant Professors (*P* = 1.34×10^−234^). For all three academic ranks, men have been publishing for a significantly longer amount of time than women (Mann–Whitney’s *U* test, Assistant Professors: *P* = 0.048, Associate Professors: *P* = 4.41×10^−8^, Professors: *P* = 9.82×10^−17^).

To remove the effect of the fact that men in our dataset have been, on average, publishing for a longer amount of time, we first computed, for each faculty member, the average number of publications per year as their number of publications divided by the time since their first publication (Table 2, Fig. S2). For all three academic ranks, men have authored a significantly higher number of publications per year than women (Mann–Whitney’s *U* test, Assistant Professors: *P* = 1.86×10^−8^, Associate Professors: *P* = 0.0004, Professors: *P* = 2.22×10^−9^). Second, given the possibility that typical publication rates and/or the productivity gap may have changed over the years, we counted, for each faculty member in our dataset, the number of publications that they authored in 2023 (Table S2). Men published on average more than women for all three academic ranks, even though the differences were only significant among Assistant Professors (Mann–Whitney’s *U* test, Assistant Professors: *P* = 0.002, Associate Professors: *P* = 0.931, Professors: *P* = 0.810). Third, we compared the average number of publications of women and men within each cohort (each cohort consisted of all faculty members who started publishing in the same year). Men authored on average more publications than women for almost every cohort, an effect that was more apparent for cohorts that started publishing earlier (Fig. 3).

**Figure 3:**
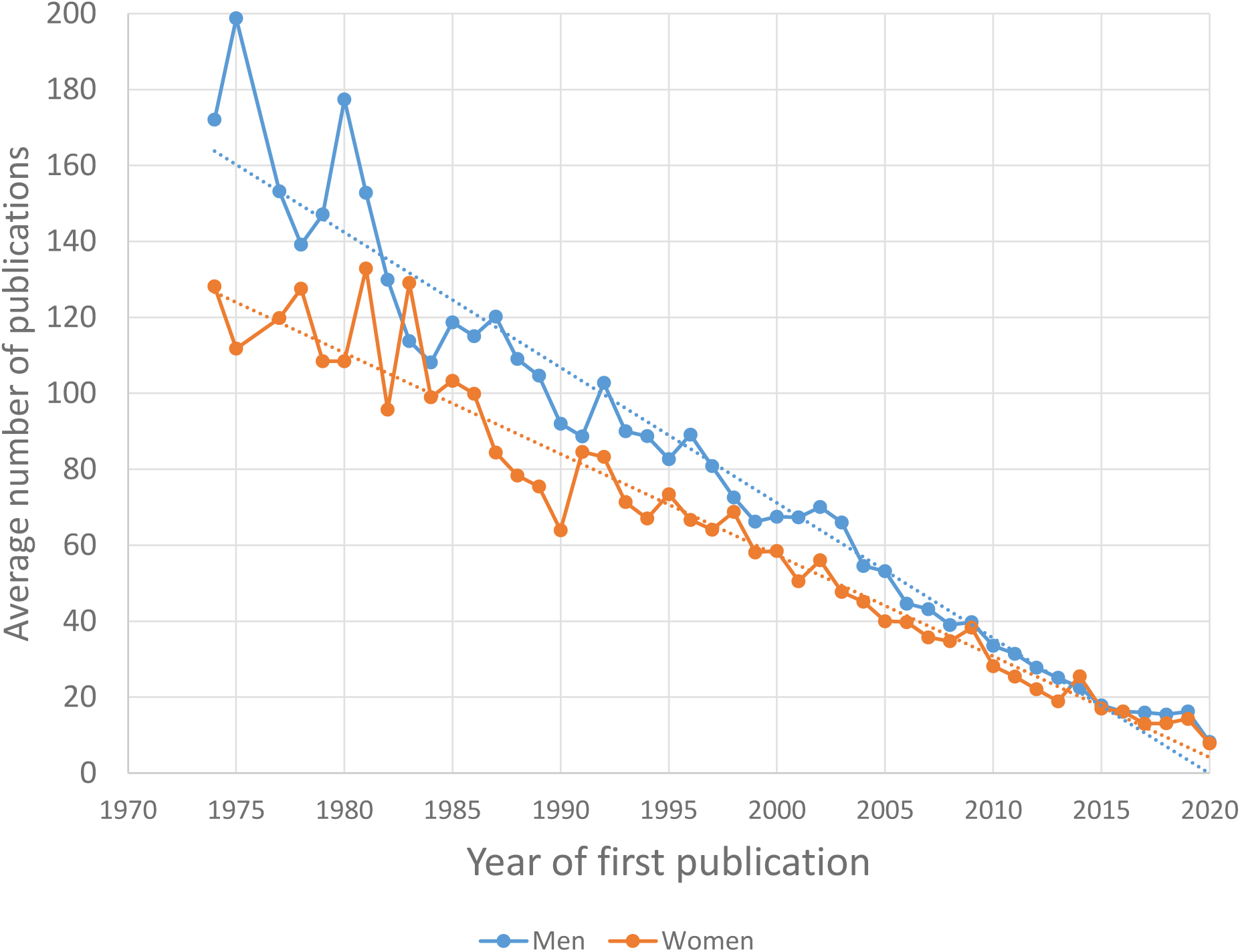
A**v**erage **number of total publications of women and men in each cohort.** Each cohort is composed of all faculty members that started publishing in a given year. Only cohorts with at least 6 women and 6 men are represented. Dotted lines represent regression lines.

**Table 2:**
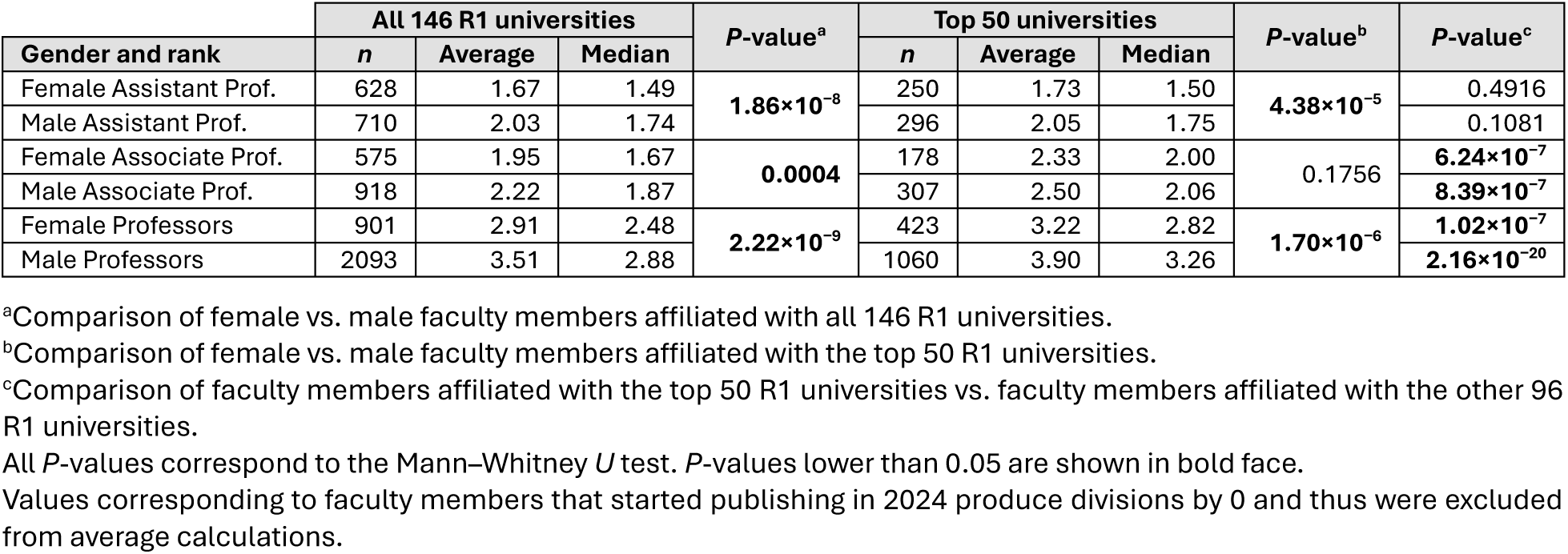
Number of publications per year.

To account for the effect of the different confounding variables simultaneously, we conducted an ANCOVA using the number of publications as the dependent variable and gender, years since first publication and ranking of the university as independent variables. The analysis showed that all independent variables had a significant effect on the number of publications (*P* ≤ 9.03×10^−14^). A similar ANCOVA using the number of publications in 2023 as the dependent variable produced similar results (*P* ≤ 0.025 for all independent variables).

Taken together, results presented in this section indicate that, among tenured and tenure-eligible Biology faculty members working at United States R1 universities, men publish at higher rates than women, and that these differences cannot be entirely attributed to differences in the amount of time that they have been publishing for or the ranking of the universities they are affiliated with.

### Male faculty members are more cited than female faculty members

For each of the faculty members in our dataset, we retrieved their total number of citations (Table S3, Fig. S3) and their *h*-index (Table 3, Fig. 4) from the Scopus database. As expected, both metrics were higher on average for Professors than for Associate Professors (Mann–Whitney *U* test, number of citations: *P* = 9.65×10^−201^, *h*-index: *P* = 1.83×10^−271^) and for Associate Professors than for Assistant Professors (number of citations: *P* = 5.01×10^−108^, *h*-index: *P* = 4.18×10^−135^). Within each rank (Assistant Professor, Associate Professor and Professor), men have on average a higher number of citations (Mann–Whitney *U* test, Assistant Professors: *P* = 2.76×10^−12^, Associate Professors: *P* = 9.82×10^−16^, Professors: *P* = 5.02×10^−18^) and a higher *h*-index (Assistant Professors: *P* = 4.66×10^−14^, Associate Professors: *P* = 1.15×10^−19^, Professors: *P* = 3.13×10^−21^) than women.

**Figure 4:**
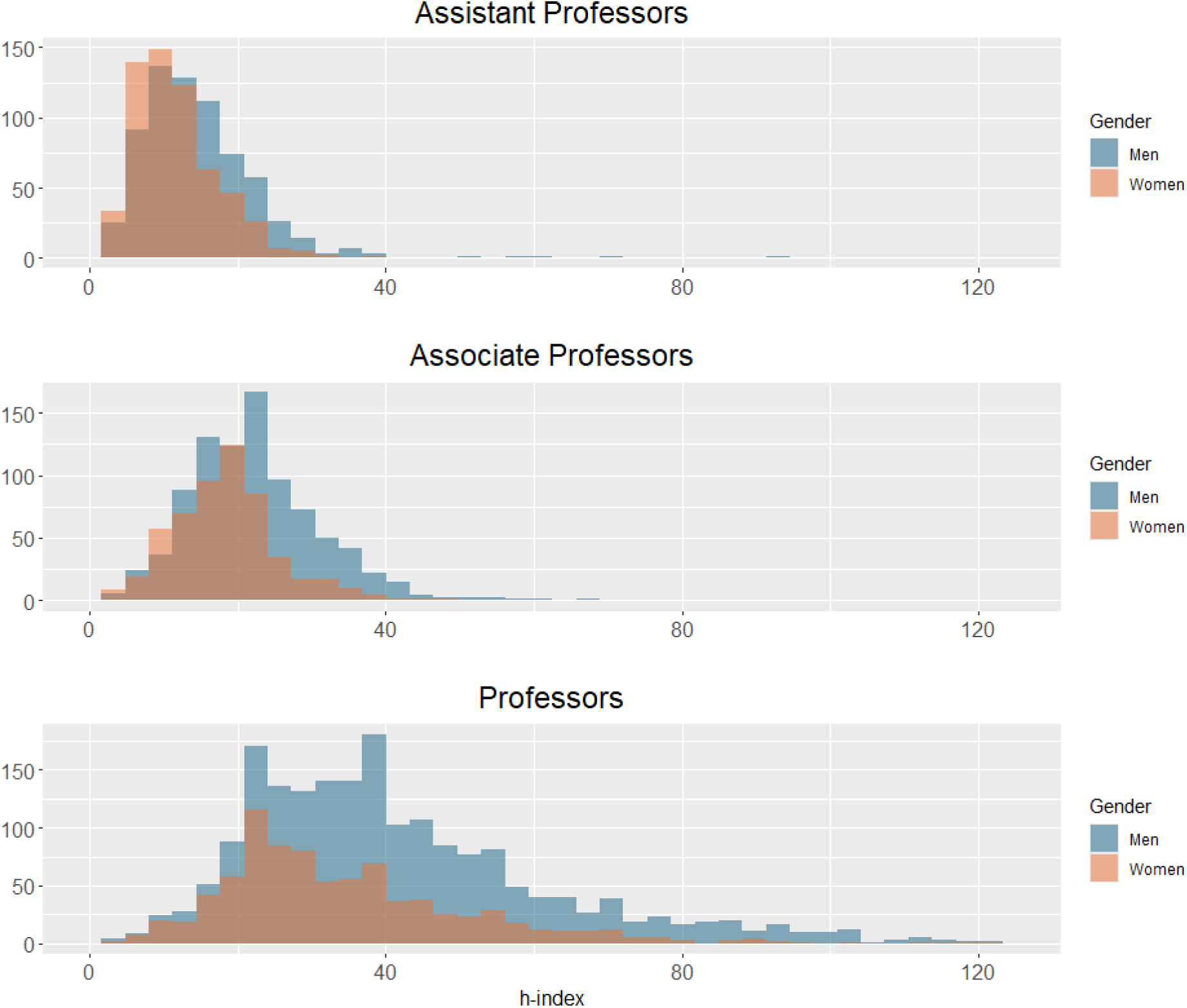
h-index of female and male faculty members of different ranks.

**Table 3:** h-index.

| Gender and rank | All 146 R1 universities |  |  | *P*-value<sup>a</sup> | Top 50 universities |  |  | *P*-value<sup>b</sup> | *P*-value<sup>c</sup> |
| --- | --- | --- | --- | --- | --- | --- | --- | --- | --- |
|  | *n* | Average | Median |  | *n* | Average | Median |  |  |
| Female Assistant Prof. | 628 | 11.61 | 11 | **4.66×10<sup>-14</sup>** | 250 | 12.54 | 12 | **7.44×10<sup>-6</sup>** | **0.0006** |
| Male Assistant Prof. | 710 | 14.58 | 13 |  | 296 | 15.15 | 14 |  | **0.0118** |
| Female Associate Prof. | 575 | 17.99 | 18 | **1.15×10<sup>-19</sup>** | 178 | 20.65 | 20 | **1.04×10<sup>-6</sup>** | **2.95×10<sup>-9</sup>** |
| Male Associate Prof. | 918 | 22.28 | 21 |  | 307 | 24.57 | 23 |  | **4.95×10<sup>-9</sup>** |
| Female Professors | 901 | 34.44 | 30 | **3.13×10<sup>-21</sup>** | 423 | 39.6 | 36 | **2.53×10<sup>-12</sup>** | **3.24×10<sup>-17</sup>** |
| Male Professors | 2093 | 42.92 | 38 |  | 1060 | 49.57 | 44 |  | **1.00×10<sup>-42</sup>** |

<sup>a</sup>Comparison of female vs. male faculty members affiliated with all 146 R1 universities.
<sup>b</sup>Comparison of female vs. male faculty members affiliated with the top 50 R1 universities.
<sup>c</sup>Comparison of faculty members affiliated with the top 50 R1 universities vs. faculty members affiliated with the other 96 R1 universities.
All *P*-values correspond to the Mann–Whitney *U* test. *P*-values lower than 0.05 are shown in bold face.

Are the higher citation metrics of male faculty members due to the fact that, on average, they have been publishing and being cited for a longer amount of time? To test this possibility, we first calculated, for each faculty member, the number of citations per year by dividing their total number of citations by the number of years since their first publication (Table S4). This ratio is higher for Professors than for Associate Professors (Mann–Whitney *U* test, *P* = 1.24×10^−105^) and for Associate Professors than for Assistant Professors (*P* = 1.38×10^−33^). Within each rank, men have a higher number of citations per year than women (Mann–Whitney *U* test, Assistant Professors: *P* = 2.24×10^−12^, Associate Professors: *P* = 1.46×10^−10^, Professors: *P* = 4.61×10^−11^). Second, we classified all professors into cohorts according to the year of their first publication and calculated the average number of citations of women and men within each cohort. For almost all cohorts, men exhibited a higher average number of citations (Fig. 5) and a higher average *h*-index (Fig. S4). Third, we calculated the *m*-index (Hirsch 2005) of each faculty member by dividing their *h*-index by the number of years since their first publication (Table S5, Fig. S5). On average, *m*-indexes were higher for Professors than for Associate Professors (Mann–Whitney *U* test, *P* = 1.58×10^−56^), but no significant differences were found between Associate Professors and Assistant Professors (*P* = 0.959). Within each rank, men had a significantly higher *m*-index than women (Mann–Whitney *U* test, Assistant Professors: *P* = 1.24×10^−11^, Associate Professors: *P* = 5.94×10^−6^, Professors: *P* = 1.25×10^−6^).

**Figure 5:**
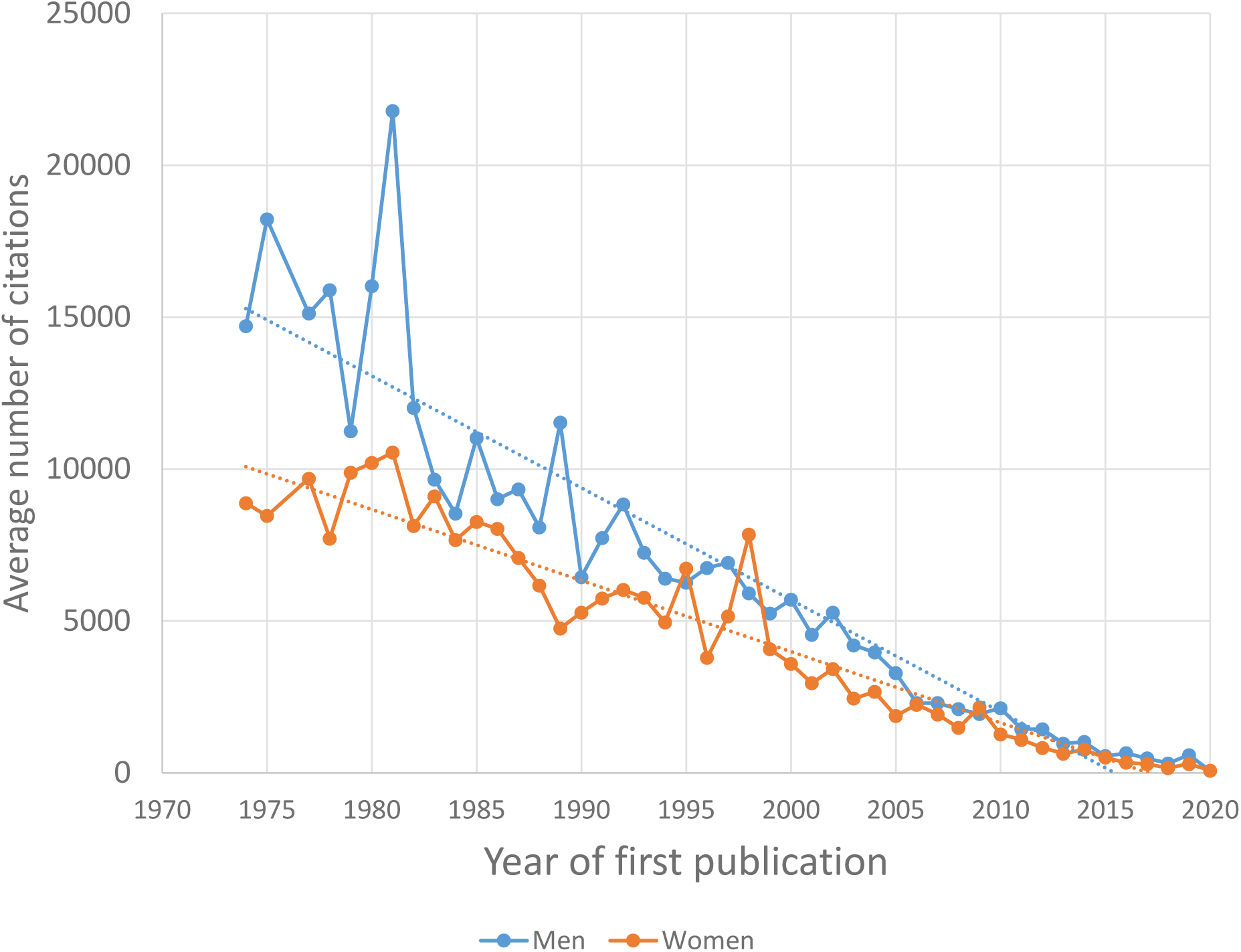
Average number of total citations of women and men in each cohort. **Each cohort is** composed of all faculty members that started publishing in a given year. Only cohorts with at least 6 women and 6 men are represented. Dotted lines represent regression lines.

Are the higher citation rates of male faculty members due to the fact that, on average, they have authored a higher number of publications? To test this possibility, we calculated, for each faculty member, the number of citations per publication by dividing their total number of citations by their total number of publications (Table S6). This ratio is higher for Professors than for Associate Professors (Mann–Whitney *U* test, *P* = 2.15×10^−37^) and for Associate Professors than for Assistant Professors (*P* = 3.69×10^−32^). Within each rank, men have a higher number of citations per publication (Mann–Whitney *U* test, Assistant Professors: *P* = 2.85×10^−6^, Associate Professors: *P* = 1.70×10^−6^, Professors: *P* = 6.27×10^−6^). In addition, we classified all professors according to their total number of publications, and, for those categories with at least 6 women and 6 men (*n* = 99), we compared the average number of citations and the average *h*-index of women vs. men. For 68 of the 99 categories, men exhibit a higher average number of citations (Fig. S6), a fraction that is significantly higher than 50% of the categories (binomial test, *P* = 0.0003). For 75 of the 99 categories, men exhibit a higher average *h*-index (Fig. S7), which also significantly exceeds 50% of the categories (binomial test, *P* = 2.77×10^−7^).

To remove the confounding effect of both the year of first publication and the total number of publications simultaneously, we conducted two additional analyses. First, for each woman with available bibliometric data in our dataset (*n* = 2013), we attempted to find a man in our dataset from the same cohort (i.e., a man who started publishing in the same year) and with the same number of publications (and thus producing the same number of publications per year). If multiple matching men were found, one was selected randomly. We could find matches for 1124 of the 2013 women. Comparing the 1124 women with the 1124 men in the matched dataset, we found that, on average, women had 2352.38 citations and a *h*-index of 19.01, whereas men had 2698.38 citations and an *h*-index of 19.96. In 604 of the 1124 woman–man pairs, the man had more citations than the woman, which significantly exceeds 50% of the pairs (binomial test, *P* = 0.013). In 586 of the 1124 pairs, the man had a higher *h*-index than the woman, which did not significantly exceed 50% of the pairs (binomial test, *P* = 0.161). Second, we conducted an ANCOVA using the *h*-index as the dependent variable and gender, years since first publication, total number of publications and ranking of the university as independent variables. The analysis showed that all independent variables had a significant effect on the *h*-index (*P* ≤ 2.93×10^−9^).

In summary, results presented in this section indicate that men tend to be more cited than women, regardless of their number of publications, the length of their careers, the rate at which they publish or the ranking of their institutions.

### Male and female faculty members exhibit similar patterns of co-authorship

We considered whether the higher number of citations received by male-authored publications could be due to differences in authorship patterns. Articles with a high number of co-authors tend to be highly cited (H. Shen et al. 2021), so we tested whether male-authored publications in our dataset tend to have a higher number of co-authors. For each faculty member in our dataset, we computed the average number of authors per publication, and we calculated the median and average for faculty members in each of the six categories (female Assistant Professors, male Assistant Professors, female Associate Professors, male Associate Professors, female Professors, male Professors; Table S7). No significant differences were found between female Assistant Professors and male Assistant Professors (Mann–Whitney’s *U* test, *P* = 0.170), between female Associate Professors and male Associate Professors (*P* = 0.983), or between female Professors and male Professors (*P* = 0.109), indicating that the higher number of citations received by male-authored publications is not due to them having more co-authors. Nonetheless, as expected from their higher number of publications, men tend to have larger co-author networks (i.e., a higher number of researchers with whom they have co-authored publications at some point of their careers) than women in the same rank (Table 4).

**Table 4:** Size of coauthor network.

| Gender and rank | All 146 R1 universities |  |  | *P*-value<sup>a</sup> | Top 50 universities |  |  | *P*-value<sup>b</sup> | *P*-value<sup>c</sup> |
| --- | --- | --- | --- | --- | --- | --- | --- | --- | --- |
|  | *n* | Average | Median |  | *n* | Average | Median |  |  |
| Female Assistant Prof. | 628 | 119.20 | 75.0 | **1.59×10<sup>-5</sup>** | 250 | 129.75 | 86.0 | **0.0066** | **0.0042** |
| Male Assistant Prof. | 710 | 169.92 | 93.0 |  | 296 | 168.13 | 100.0 |  | **0.0064** |
| Female Associate Prof. | 575 | 160.58 | 106.0 | **6.70×10<sup>-5</sup>** | 178 | 213.46 | 140.0 | 0.0571 | **1.52×10<sup>-7</sup>** |
| Male Associate Prof. | 918 | 212.13 | 130.0 |  | 307 | 258.47 | 160.0 |  | **1.22×10<sup>-9</sup>** |
| Female Professors | 901 | 318.99 | 198.5 | **1.03×10<sup>-6</sup>** | 423 | 369.88 | 233.5 | **0.0008** | **5.03×10<sup>-9</sup>** |
| Male Professors | 2093 | 389.68 | 241.0 |  | 1060 | 457.42 | 286.0 |  | **4.50×10<sup>-18</sup>** |

<sup>a</sup>Comparison of female vs. male faculty members affiliated with all 146 R1 universities.
<sup>b</sup>Comparison of female vs. male faculty members affiliated with the top 50 R1 universities.
<sup>c</sup>Comparison of faculty members affiliated with the top 50 R1 universities vs. faculty members affiliated with the other 96 R1 universities.
All *P*-values correspond to the Mann–Whitney *U* test. *P*-values lower than 0.05 are shown in bold face.

For each faculty member, we computed the percent of publications as first author, as last author, and as single author. In general, these quantities did not significantly differ between men and women; however, we found the following significant differences: (1) among Assistant Professors, women exhibit a higher percentage of publications as first authors than men (Mann–Whitney’s *U* test, *P* = 6.73×10^−6^; Table S8); (2) among Associate Professors, women exhibit a higher percentage of publications as last authors than men (*P* = 0.019; Table S9); and (3) among Professors, men exhibit a higher percentage of publications as single authors than men (*P* = 0.0007; Table S10).

### Male faculty members tend to publish in journals with higher impact factors

For each faculty member in our dataset, we computed the average journal impact factor across all their publications, and we calculated the median and average for faculty members in each of the six categories (female Assistant Professors, male Assistant Professors, female Associate Professors, male Associate Professors, female Professors, male Professors; Table S11). Only journals whose 2023 impact factors could be found in the 2023 *Journal Citation Reports* were used for average calculations. Average impact factors were significantly higher for Assistant Professors than for Professors (Mann–Whitney’s *U* test, *P* = 0.013), and significantly higher for Professors than for Associate Professors (*P* = 8.18×10^−7^). Within each of the three academic ranks, average impact factors were significantly higher for men than for women (Mann–Whitney’s *U* test, Assistant Professors: *P* = 2.39×10^−7^, Associate Professors: *P* = 0.0007, Professors: *P* = 0.015).

We considered the possibility that men’s tendency to publish in higher-impact factor journals could explain their higher citation rates. However, an ANCOVA using the *h*-index as the dependent variable and gender, years since first publication, total number of publications, ranking of the university and average impact factor as independent variables showed that all independent variables had a significant effect on the *h*-index (*P* ≤ 8.19×10^−6^), indicating that women in our dataset are less cited than men in our dataset regardless of the impact factors of the journals in which they publish.

### Publication and citation patterns of faculty at the top 50 United States universities

We recalculated all averages and medians for faculty members in the six different categories (female Assistant Professors, male Assistant Professors, female Associate Professors, male Associate Professors, female Professors, male Professors) at the top 50 United States universities. We found that, for the four most senior categories (female Associate Professors, male Associate Professors, female Professors and male Professors), faculty at the top 50 universities had authored a significantly higher number of publications (Mann–Whitney’s *U* test, *P* ≤ 1.19×10^−5^; Table 1) and more publications per year (*P* ≤ 8.39×10^−7^; Table 2) than faculty at the other R1 universities. For all six categories, faculty members at the top 50 universities had higher numbers of publications in 2023 (*P* ≤ 0.028; Table S2), higher numbers of citations (*P* ≤ 3.35×10^−5^; Table S3), higher *h*-indexes (*P* ≤ 0.012; Table 3), higher numbers of citations per year (*P* ≤ 2.21×10^−5^; Table S4), higher *m*-indexes (*P* ≤ 0.013; Table S5), higher numbers of citations per publication (*P* ≤ 1.11×10^−5^; Table S6), higher numbers of co-authors per publication (*P* ≤ 0.038; Table S7), larger co-author networks (*P* ≤ 0.006; Table 4), smaller percentages of publications as first authors (*P* ≤ 0.005; Table S8), and a higher average impact factor (P ≤ 2.54×10^−15^; Table S11). For four of the categories (female Assistant Professors, female Associate Professors, female Professors and male Professors), faculty at the top 50 universities exhibited a higher percentage of publications as last author (Mann–Whitney’s *U* test, *P* ≤ 0.024; Table S9). Among male Associate Professors, faculty at the top 50 universities had been publishing for fewer years (*P* = 0.008; Table S1) and exhibited smaller percentages of publications as single authors (*P* = 0.029; Table S10). Among male Professors, faculty at the top 50 universities had been publishing for longer (*P* = 0.011; Table S1). Among female Professors, faculty at the top 50 universities exhibited higher percentages of publications as single authors (*P* = 0.001; Table S10).

In summary, results presented in this section indicate that, regardless of their gender and their academic rank, faculty members at the top 50 United States universities publish at higher rates and in higher-impact factor journals than faculty at the other R1 universities. In addition, they tend to receive more citations, regardless of their number of publications, their publication rates, and the length of their careers.

### Female and male faculty members require the same amount of time to attain the rank of Professor

We first considered whether men and women in our dataset had been promoted to the rank of Professor at different speeds. For each cohort (each cohort consisted of all faculty members who started publishing in the same year) with at least 10 female and 10 male faculty members, we computed the percentage of Professors (i.e., the percent of the faculty members that have attained the rank of Professor) separately among women and men. The percentage of Professors increased over time describing similar asymptotic curves for women and for men (Fig. 6). For 22 out of the 41 cohorts, the percentage of Professors was higher among women than among men, a number that was not significantly different from 50% (binomial test, *P* = 0.755). These results indicate that women and men in our dataset were promoted to the rank of Professor at the same speeds. For each woman with available bibliometric data in our dataset (*n* = 2013), we found a man in our dataset from the same cohort (if multiple matching men were found, one was selected randomly), resulting in 2013 woman–man pairs. The 2013 women in these pairs included 604 Assistant Professors, 557 Associate Professors and 852 Professors. The 2013 men included 614 Assistant Professors, 564 Associate Professors and 835 Professors. The fraction of Professors was similar among women (42.32%) and among men (41.48%; Fisher’s exact test, *P* = 0.609), confirming that women and men in our dataset attained the rank of Professor in a similar timeframe.

**Figure 6.**
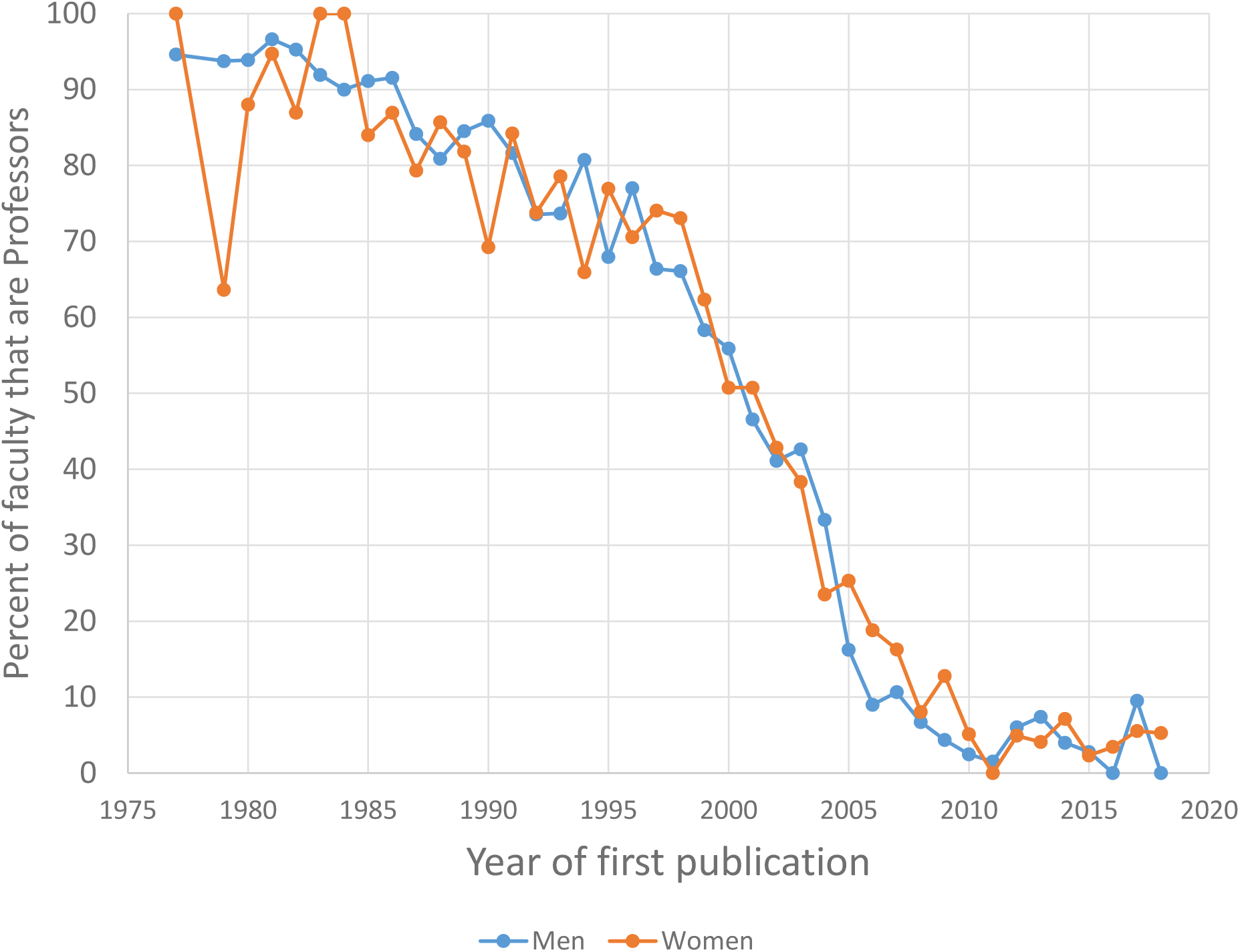
Percentage of tenured and tenure-eligible faculty members that have attained the rank of Professor among women and men in each cohort. Each cohort is composed of all faculty members that started publishing in a given year. Only cohorts with at least 10 women and 10 men are represented.

Second, we considered whether women and men required a different number of publications to achieve the rank of Professor. We classified faculty members into groups according to their number of publications. For each group with at least 10 female and 10 male faculty members, we computed the percentage of Professors among women and among men (Fig. S8). In 34 out of the 66 groups, women were more likely to be Professors than men, which did not significantly differ from 50% of the groups (binomial test, *P* = 0.902). These results indicate that women and men in our datasets required roughly the same number of publications to attain the rank of Professor. For each woman with available bibliometric data in our dataset (*n* = 2013), we attempted to find a man in our dataset with the same number of publications. We could find matches for 1997 of the 2013 women. The 1997 women in these pairs included 604 Assistant Professors, 557 Associate Professors and 836 Professors. The 1997 men included 613 Assistant Professors, 598 Associate Professors and 786 Professors. The fraction of Professors was similar among women (41.86%) and among men (39.36%; Fisher’s exact test, *P* = 0.114), confirming that women and men require a similar number of publications to attain the rank of Professor.

Third, we considered whether women and men required a different *h*-index to be promoted to the rank of Professor. We classified faculty members according to their *h*-index. For each group with at least 10 men and 10 women, we calculated the percent of Professors among men and among women separately (Fig. S9). For 30 out the 48 categories, the percentage of Professors was higher among women than among men, a number that did not significantly differ from 50% of the groups (binomial test, *P* = 0.111). These results indicate that women and men in our dataset required similar *h*-indexes to attain the rank of Professor. For each woman with available bibliometric data in our dataset (*n* = 2013), we found a man in our dataset with the same *h*-index. The 2013 women in these pairs included 604 Assistant Professors, 557 Associate Professors and 852 Professors. The 2013 men included 600 Assistant Professors, 565 Associate Professors and 848 Professors. The fraction of Professors was similar among women (42.32%) and among men (42.13%; Fisher’s exact test, *P* = 0.924), confirming that women and men require similar *h*-indexes to attain the rank of Professor.

Last, to control for the number of years since first publication and the number of publications simultaneously, we used the 1124 woman–man pairs described above (each pair was composed of a female and a male faculty member who started publishing in the same year and had the same number of publications). The 1124 women in these pairs included 427 Assistant Professors, 337 Associate Professors and 360 Professors. The 1124 men included 458 Assistant Professors, 354 Associate Professors and 312 Professors. The fraction of Professors was significantly higher among women (32.03%) than among men (27.76%; Fisher’s exact test, *P* = 0.030), indicating that women in our dataset required fewer publications per year than their male peers to attain the rank of Professor.

## DISCUSSION

### The gender representation gap

Previous work has shown that women are underrepresented in many academic fields, including STEMM fields, and including Biology (Baker and Koedel 2024; Ceci et al. 2009, 2014; Holman et al. 2018; Hornig 1980; Rushworth et al. 2021). For instance, Holman et al. (Holman et al. 2018) found that women account for 38% of authors of research articles published in Biology journals indexed in PubMed. It should be noted, however, that estimates of female representation based on analyses of published articles reflect past rather than current trends, and depend not only on how many female researchers work (or have worked) on a given field, but also on their relative publication rates compared to male researchers. In addition, the authors of research articles include not only tenured and tenure-eligible faculty, but also trainees and other kinds of professionals. Nonetheless, our sample of 5825 active Biology faculty members includes 2104 women, a fraction (36.12%) that is very similar to the one estimated by Holman et al. (Holman et al. 2018). Studies focusing on ecologists and evolutionary biologists also produced similar estimates of female representation: women account for 28–31% of the authors, 29–32% of editors and 27% of reviewers of articles published in Ecology and Evolution academic journals (Alvarez-Ponce and Vesper 2025; Edwards et al. 2018; Fox et al. 2019).

In agreement with previous findings that women are particularly underrepresented at the top academic ranks (Ceci et al. 2009, 2014; Ginther and Kahn 2004; Holman et al. 2018; Hornig 1980; Rushworth et al. 2021), women represent 46.94% of Assistant Professors, 38.51% of Associate Professors and 30.09% of Professors in our dataset (Fig. 1). As mentioned above, this could be due to the steady increase in the number of women entering the field, to female biologists abandoning the field at higher rates than their male peers, or to women being promoted more slowly than men (Alvarez-Ponce and Vesper 2025; Ceci et al. 2014; Fox et al. 2019; Goulden et al. 2011; Holman et al. 2018; Shaw and Stanton 2012).

Previous studies have shown that women are particularly underrepresented at top research-producing institutions (Ceci et al. 2009; Hornig 1980). In contrast, we found a weak but significantly positive correlation between universities’ ranking and the fraction of female faculty members at their Biology departments (i.e., Biology departments at higher-ranking universities tend to employ a higher percent of women; ρ = 0.172, *n* = 146, *P* = 0.038). In addition, the percent of faculty that are female is only slightly lower in Biology departments at the top 50 universities (33.85%) than in all R1 universities combined (36.12%). It should be noted, however, that our analysis only included departments at R1 universities (i.e., top research-intensive universities), and that it is possible that female representation is higher at the Biology departments of other kinds of institution.

### The gender productivity gap

On average, female researchers appear to publish less than male researchers in most fields (Astegiano et al. 2019; J. R. Cole 2024; Holman et al. 2018; Larivière et al. 2013; Martin 2012; West et al. 2013), but the trend seems to be absent, and even reversed, in certain fields and social contexts (Arensbergen et al. 2012; Duch et al. 2012; Frandsen et al. 2020; Schucan Bird 2011). Thus, it is essential to evaluate the trend separately for each field and context. A number of studies have shown that female researchers publish on average fewer papers per year than their male peers (Duch et al. 2012; Symonds et al. 2006), whereas others suggest that they publish at similar rates as their male peers, but for a shorter period of time (i.e., they tend to have shorter careers; Huang et al. 2020). While some studies have analyzed the publication records of all researchers that have published in a certain area at some point (e.g., Huang et al. 2020), we focused on a group of researchers that have secured tenure or a tenure-track appointment at an R1 university. As a result, we are focusing on a group of researchers that are especially successful and that have had relatively long careers (Fig. S1, Table S1).

We provide unequivocal evidence that, among Biology professors affiliated with R1 universities, men produce a higher number of publications per year than their female peers. The median number of publications per year is 11.98–16.78% higher for men than for women, depending on the academic rank (Fig. S2; Table 2), and differences remain significant after controlling for university ranking. In addition, the total number of publications increases faster over men’s careers than over women’s careers (Fig. 3). Finally, ANCOVA analyses showed that male authors had produced more publications (both throughout their entire careers and in 2003), even after controlling for career length and university ranking.

One of the reasons that has been invoked to explain women’s lower publication rates is the fact that they are less likely to secure research funding than their male peers. In agreement with that hypothesis, Duch et al. (Duch et al. 2012) found that the productivity gap is more pronounced in those fields that require more resources to produce research. On average, biological research is expensive, even though research costs are lower in certain subfields, e.g. those that are more theoretical and/or computational.

Another potential reason for the productivity gap is that female-authored work may be subjected to higher standards by editors and/or referees. This hypothesis has been supported by a number of analyses of published articles, especially in Economics journals. First, Card et al. (Card et al. 2020) showed that female-authored Economics articles tend to be more cited than comparable male-authored articles published in the same journals, which could be due to them being of higher quality. Second, Hengel (Hengel 2022) showed that female economists tend to rewrite their abstracts to increase their readability more substantially than their male peers. Third, a handful of analyses have shown that female-authored manuscripts spend longer under review than comparable male-authored manuscripts (Alexander et al. 2023; Argilés-Bosch et al. 2025; Bruns et al. 2025; Hagan et al. 2020; Hengel 2022; Tutuncu and Dag 2024). However, this is not the case in all fields: in Evolutionary Biology, female-and male-authored manuscripts spend similar amounts of time under review once confounding factors are controlled for (Alvarez-Ponce and Vesper 2025), and in Biology as a whole, female-authored articles tend to experience *shorter* review times than male-authored articles (Alvarez-Ponce, Batz and Ramirez Torres, unpublished results).

As mentioned above, other possible explanations for the productivity gap include female academics being responsible for a disproportionate amount of childcare, domestic, teaching and/or service duties (Awung and Dorasamy 2015; Eagly 2020; Guarino and Borden 2017; Ledin et al. 2007; O’Meara et al. 2017; Rhoads and Rhoads 2012), and female academics being on average more perfectionistic, risk-averse and prone to self-criticism and self-doubt (Born et al. 2022; Coffman 2014; Croson and Gneezy 2009; Exley and Kessler 2022; Kessler et al. 2014; Möbius et al. 2022; Shastry and Shurchkov 2024)—which may be the result, at least in part, of the way in which they are treated by academia (e.g., Hengel 2022).

### The gender citation gap

A number of studies have shown that male-authored research articles tend to be more cited than comparable female-authored articles, an effect that cannot be entirely explained by self-citations or other confounding factors such as year of publication, journal, article focus, theoretical perspective, methodology, career stage, or institution (Ceci et al. 2014; Chatterjee and Werner 2021; Davenport and Snyder 1995; Ferber and Brün 2011; Larivière et al. 2013; Maliniak et al. 2013; Teich et al. 2022). Other studies, however, suggest that female-and male-authored articles are cited equally, and that the only reason men tend to be more cited is their higher rates of publication (Aksnes et al. 2011; Huang et al. 2020; Pautasso 2013). Yet another group of studies suggests that, in certain fields, female-authored articles are more cited than male-authored articles, which has been attributed to their higher quality (Card et al. 2020; Duch et al. 2012; Long 1992; Symonds et al. 2006). Again, such mixed results indicate that whether or not authors of one gender are favored over authors of the other in the citation arena depends on the specific social context.

Our analyses clearly show that, among Biology faculty members affiliated with R1 universities, men on average have higher citation metrics than their female peers, and that differences are not entirely due to men having longer or more productive careers (Figs. 4, 5 and S3– S7; Tables 3 and S3–S6). Indeed, (1) men receive, on average, a higher number of citations per year than women in the same academic rank (Table S4); (2) male-authored publications receive, on average, 14.18–19.37% more citations than publications authored by women in the same academic rank (Table S6); (3) men have, on average, higher *h*-indexes and *m*-indexes than women in the same academic rank (Figs. 4 and S5; Tables 3 and S5); (4) both the number of citations and the *h*-index increase faster over men’s careers than over women’s careers (Figs. 5 and S4) and as they accumulate more publications (Figs. S6 and S7); (5) when compared with randomly selected men from the same cohort and with the same number of publications (and thus, producing the same number of publications per year), women have on average 14.71% fewer citations and *h*-indexes that are lower by almost 1 unit; and (6) ANCOVA analyses show that men tend to have a higher *h*-index than women, even after controlling for career length, number of publications and university rank.

Research articles with many co-authors tend to be highly cited (Bosquet and Combes 2013; H. Shen et al. 2021; Thelwall et al. 2023), and it has been suggested that male researchers tend to engage in more collaborative work in certain contexts (e.g., Kyvik and Teigen 1996; Lee and Bozeman 2005, but see Abramo et al. 2013; Bozeman and Gaughan 2011; Campbell et al. 2018; Morello et al. 2023). If both trends apply to our dataset, their combination could explain, at least in part, why male-authored publications tend to be more cited. We did find a positive correlation between publications’ number of authors and number of citations for all years since 1983 (e.g., ρ = 0.269, *n* = 13,701, *P* = 2.81×10^−226^ for 2023; for prior years, the number of publications in our dataset was lower than 1050, limiting the statistical power of our tests). However, we found no differences in men’s vs. women’s average number of co-authors per publication (Table S7), indicating that the reason men in our dataset are more cited is not that their publications tend to have more co-authors.

It has been shown that men are more likely than women to publish in high-impact factor journals (Bendels et al. 2018; Krishnamurthy et al. 2017; Y. A. Shen et al. 2018), mostly because women are less likely to submit their work to such journals to avoid rejection (Basson et al. 2023). In agreement with these results, we show that the average impact factor of male-authored articles is significantly higher than that of female-authored articles (Table S11). The trend is especially pronounced among Assistant Professors and among Associate Professors, which could be due to the fact that impact factors have become increasingly relevant over the last decades (Garfield 1999). Since articles published in high-profile journals tend to enjoy high visibility, we considered the possibility that the trend could explain, at least in part, why male-authored publications tend to be more cited. However, our multivariate analyses show that gender has an effect on the *h*-index, even after controlling for years since first publication, total number of publications, ranking of the university and average impact factor, indicating that journal impact factors do not totally explain why male academics tend to be more cited.

It has also been shown that male academics tend to have larger, largely male collaborator networks (Ductor et al. 2023; Mcdowell and Smith 1992; Rothstein and Davey 1995), which may contribute to their higher citation rates (Bosquet and Combes 2013). Among other reasons, researchers tend to be familiar with and interested in the work of their collaborators, and are therefore likely to cite them. Even though male-authored publications in our dataset do not tend to have more co-authors (Table S7), men in our dataset tend to have larger co-author networks (i.e., a higher number of researchers with whom they have co-authored publications at some point; Table 4) thanks to their longer publication records (Table 1). Thus, male-authored publications in our dataset may be more cited, at least in part, due to male authors having larger collaborator networks.

We cannot discard other previously proposed explanations, such as: (1) researchers being more prone to cite the most productive authors because of their reputation (the so-called “Mathew effect”; Merton 1968) or because they are more likely to be familiar with them (the so-called “fast-food effect”; Symonds et al. 2006); (2) more productive researchers being more likely to produce highly-cited publications due to a “lottery effect” (Kelly and Jennions 2006); or (3) male researchers being more likely to self-cite, which increases not only their citation metrics, but also their visibility, which in turn increases their likelihood of being cited by other researchers (Fowler and Aksnes 2007).

### No apparent gender promotion gap

A number of studies have shown that female academics are less likely to obtain tenure than their male peers and that, when they do, they take longer to achieve it and to be promoted to the rank of Professor (Ginther and Kahn 2004; Heijstra et al. 2015; Long et al. 1993; Weisshaar 2017). Other studies, however, have found that men and women move up the academic ladder at a similar pace (Ginther and Kahn 2019).

We found that male and female Professors in our dataset attained their current rank at similar speeds: as time since first publication increases, the fraction of faculty members that are full professors increases asymptotically, describing similar trajectories for female and male faculty members (Fig. 6). In addition, when women were compared with randomly selected men from the same cohort, both men and women were equally likely to be Professors. This indicates that women and men in our dataset were promoted at similar rates, despite women publishing and being cited at lower rates. Indeed, when women were compared with randomly selected men from the same cohort and with the same number of publications (and thus producing the same amount of publications per year), women were slightly but significantly more likely than men to be Professors.

This should not be interpreted as evidence that female faculty members are promoted with less merit, given that women may require investing a higher amount of effort per publication than their male peers due to lower resource availability and to how they are treated by academia (e.g., Card et al. 2020; Eagly 2020; Hengel 2022). In addition academics’ contributions to their fields, communities and institutions go far beyond academic publications, and women often take on greater teaching and/or service responsibilities than their male peers (Eagly 2020; Guarino and Borden 2017; O’Meara et al. 2017), which are also important factors for promotion (Sabatier et al. 2006).

It should be noted that our dataset only includes tenured and tenure-eligible faculty members at R1 universities. Thus, it excludes individuals that have left academia or moved to other kinds of academic institutions, which is likely to occur when prospects of tenure or promotion are uncertain, or when tenure or promotion are denied. Female academics are more likely than their male peers to leave academia (Goulden et al. 2011; Shaw and Stanton 2012), but our dataset only includes those individuals that did not leave academia—presumably because they anticipated/anticipate earning tenure and promotion. Our results might have been different if we had analyzed a cohort of scientists that received PhDs or that were hired as Associate Professors during a certain time period, as some of the previous studies of promotion rates have done (Ginther and Kahn 2004, 2019; Long et al. 1993; Weisshaar 2017).

### Limitations of our study

Our analyses rely on automatically generated Scopus bibliometric profiles, and thus a certain error rate is unavoidable. However, it should be noted that: (1) both Biomedicine and the Natural Sciences are well represented in Scopus (Baas et al. 2020; Mongeon and Paul-Hus 2016); (2) comparison of manually-curated datasets has shown that Scopus has very high recall and precision rates (94.4% and 98.1%, respectively, according to a report published in 2020; Baas et al. 2020); (3) compared with the other leading bibliometric database—Web of Science—Scopus covers a wider range of publications and citations (Visser et al. 2021; Waltman 2016); (4) compared with Google Scholar, Scopus is more accurate and reliable—for instance, Google Scholar often indexes non-academic documents and items that are not stand-alone publications (Orduna-Malea et al. 2017)—; and (5) many academics do not have a Google Scholar profile, especially those with lower citation metrics (Kim and Grofman 2020).

Both databases are known to suffer from a number of limitations (for a review, see Pranckutė 2021). However, such limitations are expected to have only a moderate impact on our results. First, literature written in languages other than English is underrepresented in both databases (even though it is better represented in Scopus; Vera-Baceta et al. 2019); however, biologists, and especially those based in the United States, tend to publish most of their results in English. Second, books are also underrepresented in both databases compared with research journals (Waltman 2016); however, biologists publish most of their primary literature in research journals. Third, whereas Scopus-indexed publications date back to 1788, their references only go back to the 1970s (https://www.elsevier.com/products/scopus/content; last accessed July 8, 2025); however, only 0.088% of the publications in our dataset date from before 1970, and the issue is expected to have only a minor effect on the citation metrics of the oldest faculty members in our dataset (those that were over ∼80 years old at the time of data collection—a very small fraction considering that excluded emeritus faculty from our dataset).

Bibliometric comparisons of male vs. female academics can be somewhat affected by the fact that women are more likely than men to change their names—in many countries, including the United States, women often adopt their spouse’s last name when they get married. This can result in them having multiple Scopus profiles (Pellack and Kappmeyer 2011), which would impact their cumulative bibliometric indicators (including their total number of publications, total number of citations, and *h*-index). Nonetheless, female academics are less likely than the average woman to change their names (Goldin and Shim 2004), and when they do, that only moderately impacts their cumulative bibliometric indicators (Holliday et al. 2014). In addition, normalized indicators (including the number of publications per year, the number of publications in 2023, the number of citations per year, the number of citations per publication, and the *m*-index) will not be affected by a name change, or not to a great extent.

## ACKNOWLEDGEMENTS

The authors are grateful to Niveda Iyengar for assistance with data collection.

## AUTHOR CONTRIBUTIONS

DAP conceived and supervised the study, assisted in data collection, analyzed the data and wrote the manuscript. JV was the main responsible for data collection.

## ETHICS DECLARATION

Prior to data collection, the University of Nevada, Reno’s Institutional Review Board determined that the project did not involve human research since it was entirely based on publicly available data and thus did not require institutional approval. We do not provide any personally identifiable information.

