## Supplementary Tables and Figures for "The publication and citation gender gap in Biology: bibliometric analysis of Biology faculty at top United States universities"

### Table S1: Years since first publication

| | All 146 R1 universities | | | *P*-value[a] | Top 50 universities | | | *P*-value[b] | *P*-value[c] |
|---|---|---|---|---|---|---|---|---|---|
| **Gender and rank** | ***n*** | **Average** | **Median** | | ***n*** | **Average** | **Median** | | |
| Female Assistant Prof. | 628 | 13.78 | 13.5 | **0.0476** | 250 | 14.04 | 14.0 | 0.6409 | 0.2587 |
| Male Assistant Prof. | 710 | 14.35 | 14.0 | | 296 | 14.35 | 14.0 | | 0.7083 |
| Female Associate Prof. | 575 | 21.68 | 20.0 | **$4.41\times10^{-8}$** | 178 | 21.43 | 20.0 | **0.0233** | 0.6819 |
| Male Associate Prof. | 918 | 23.86 | 23.0 | | 307 | 22.89 | 21.0 | | **0.0083** |
| Female Professors | 901 | 30.44 | 29.0 | **$9.82\times10^{-17}$** | 423 | 31.09 | 30.5 | **$4.20\times10^{-8}$** | 0.1085 |
| Male Professors | 2093 | 33.77 | 33.0 | | 1060 | 34.37 | 34.0 | | **0.0107** |

[a]*P*-values corresponding to the Mann-Whitney *U* test comparing female vs. male academics affiliated with all 146 R1 universities.
[b]*P*-values corresponding to the Mann-Whitney *U* test comparing female vs. male academics affiliated with the 50 top R1 universities.
*P*-values lower than 0.05 are shown in bold face.

### Table S2: Number of publications in 2023

| | All 146 R1 universities | | | *P*-value[a] | Top 50 universities | | | *P*-value[b] | *P*-value[c] |
|---|---|---|---|---|---|---|---|---|---|
| **Gender and rank** | ***n*** | **Average** | **Median** | | ***n*** | **Average** | **Median** | | |
| Female Assistant Prof. | 628 | 2.23 | 2 | **0.0017** | 250 | 2.51 | 2 | 0.0912 | **0.0129** |
| Male Assistant Prof. | 710 | 2.67 | 2 | | 296 | 2.85 | 2 | | **0.0276** |
| Female Associate Prof. | 575 | 2.54 | 2 | 0.9315 | 178 | 3.35 | 2 | 0.8931 | **$6.37\times10^{-5}$** |
| Male Associate Prof. | 918 | 2.58 | 2 | | 307 | 3.25 | 2 | | **$6.81\times10^{-7}$** |
| Female Professors | 901 | 3.28 | 2 | 0.8102 | 423 | 3.75 | 3 | 0.9398 | **0.0003** |
| Male Professors | 2093 | 3.58 | 2 | | 1060 | 3.88 | 3 | | **$2.55\times10^{-6}$** |

[a]Comparison of female vs. male faculty members affiliated with all 146 R1 universities.
[b]Comparison of female vs. male faculty members affiliated with the top 50 R1 universities.
[c]Comparison of faculty members affiliated with the top 50 R1 universities vs. faculty members affiliated with the other 96 R1 universities.
All *P*-values correspond to the Mann–Whitney *U* test. *P*-values lower than 0.05 are shown in bold face.

### Table S3: Number of citations

| | All 146 R1 universities | | | *P*-value[a] | Top 50 universities | | | *P*-value[b] | *P*-value[c] |
|---|---|---|---|---|---|---|---|---|---|
| **Gender and rank** | ***n*** | **Average** | **Median** | | ***n*** | **Average** | **Median** | | |
| Female Assistant Prof. | 628 | 997.8 | 600.5 | **$2.76\times10^{-12}$** | 250 | 1215.2 | 754.0 | **$2.99\times10^{-7}$** | **$3.35\times10^{-5}$** |
| Male Assistant Prof. | 710 | 1603.4 | 900.0 | | 296 | 1815.0 | 1129.0 | | **$1.48\times10^{-6}$** |
| Female Associate Prof. | 575 | 2053.9 | 1524.0 | **$9.82\times10^{-16}$** | 178 | 2796.3 | 2071.0 | **$5.79\times10^{-5}$** | **$7.38\times10^{-11}$** |
| Male Associate Prof. | 918 | 3015.4 | 2175.0 | | 307 | 3887.3 | 2689.0 | | **$3.58\times10^{-12}$** |
| Female Professors | 901 | 6614.5 | 3930.0 | **5.02e-18** | 423 | 8689.3 | 5669.5 | **$2.77\times10^{-10}$** | **$2.24\times10^{-18}$** |
| Male Professors | 2093 | 10759.7 | 5790.5 | | 1060 | 14284.1 | 7816.5 | | **$3.42\times10^{-42}$** |

[a]Comparison of female vs. male faculty members affiliated with all 146 R1 universities.
[b]Comparison of female vs. male faculty members affiliated with the top 50 R1 universities.
[c]Comparison of faculty members affiliated with the top 50 R1 universities vs. faculty members affiliated with the other 96 R1 universities.
All *P*-values correspond to the Mann–Whitney *U* test. *P*-values lower than 0.05 are shown in bold face.

### Table S4: Number of citations per year

| | All 146 R1 universities | | | *P*-value[a] | Top 50 universities | | | *P*-value[b] | *P*-value[c] |
|---|---|---|---|---|---|---|---|---|---|
| **Gender and rank** | ***n*** | **Average** | **Median** | | ***n*** | **Average** | **Median** | | |
| Female Assistant Prof. | 628 | 67.60 | 44.20 | **$2.24\times10^{-12}$** | 250 | 80.99 | 53.00 | **$6.11\times10^{-8}$** | **$2.21\times10^{-5}$** |
| Male Assistant Prof. | 710 | 104.77 | 62.60 | | 296 | 124.92 | 77.20 | | **$4.88\times10^{-8}$** |
| Female Associate Prof. | 575 | 99.05 | 70.06 | **$1.46\times10^{-10}$** | 178 | 134.42 | 101.74 | **0.0016** | **$3.40\times10^{-11}$** |
| Male Associate Prof. | 918 | 135.75 | 97.39 | | 307 | 181.04 | 124.26 | | **$1.48\times10^{-14}$** |
| Female Professors | 901 | 213.64 | 140.05 | **$4.61\times10^{-11}$** | 423 | 277.51 | 185.36 | **$4.33\times10^{-7}$** | **$3.14\times10^{-18}$** |
| Male Professors | 2093 | 303.86 | 182.08 | | 1060 | 394.67 | 237.93 | | **$1.18\times10^{-42}$** |

[a]Comparison of female vs. male faculty members affiliated with all 146 R1 universities.
[b]Comparison of female vs. male faculty members affiliated with the top 50 R1 universities.
[c]Comparison of faculty members affiliated with the top 50 R1 universities vs. faculty members affiliated with the other 96 R1 universities.
All *P*-values correspond to the Mann–Whitney *U* test. *P*-values lower than 0.05 are shown in bold face.
Values corresponding to faculty members that started publishing in 2024 produce divisions by 0 and thus were excluded from average calculations.

### Table S5: *m*-index

| | All 146 R1 universities | | | *P*-value[a] | Top 50 universities | | | *P*-value[b] | *P*-value[c] |
|---|---|---|---|---|---|---|---|---|---|
| **Gender and rank** | ***n*** | **Average** | **Median** | | ***n*** | **Average** | **Median** | | |
| Female Assistant Prof. | 628 | 0.87 | 0.83 | **$1.24\times10^{-11}$** | 250 | 0.93 | 0.88 | **2.30e-06** | **0.0133** |
| Male Assistant Prof. | 710 | 1.05 | 1.00 | | 296 | 1.11 | 1.00 | | **0.0010** |
| Female Associate Prof. | 575 | 0.90 | 0.84 | **$5.94\times10^{-6}$** | 178 | 1.05 | 1.00 | **0.0118** | **$3.48\times10^{-8}$** |
| Male Associate Prof. | 918 | 1.01 | 0.95 | | 307 | 1.15 | 1.09 | | **$8.03\times10^{-12}$** |
| Female Professors | 901 | 1.16 | 1.08 | **$1.25\times10^{-6}$** | 423 | 1.31 | 1.23 | **$2.49\times10^{-5}$** | **$2.08\times10^{-14}$** |
| Male Professors | 2093 | 1.29 | 1.18 | | 1060 | 1.46 | 1.35 | | **$2.79\times10^{-40}$** |

[a]Comparison of female vs. male faculty members affiliated with all 146 R1 universities.
[b]Comparison of female vs. male faculty members affiliated with the top 50 R1 universities.
[c]Comparison of faculty members affiliated with the top 50 R1 universities vs. faculty members affiliated with the other 96 R1 universities.
All *P*-values correspond to the Mann–Whitney *U* test. *P*-values lower than 0.05 are shown in bold face.
Values corresponding to faculty members that started publishing in 2024 produce divisions by 0 and thus were excluded from this analysis.

### Table S6: Number of citations per publication

| | All 146 R1 universities | | | *P*-value[a] | Top 50 universities | | | *P*-value[b] | *P*-value[c] |
|---|---|---|---|---|---|---|---|---|---|
| **Gender and rank** | ***n*** | **Average** | **Median** | | ***n*** | **Average** | **Median** | | |
| Female Assistant Prof. | 628 | 43.36 | 29.19 | **$2.85\times10^{-6}$** | 250 | 52.11 | 33.98 | **0.0002** | **$7.68\times10^{-6}$** |
| Male Assistant Prof. | 710 | 49.51 | 35.84 | | 296 | 60.35 | 44.21 | | **$1.77\times10^{-9}$** |
| Female Associate Prof. | 575 | 52.09 | 40.87 | **$1.70\times10^{-6}$** | 178 | 62.71 | 51.54 | **0.0046** | **$1.11\times10^{-5}$** |
| Male Associate Prof. | 918 | 62.18 | 48.35 | | 307 | 75.33 | 57.92 | | **$1.04\times10^{-9}$** |
| Female Professors | 901 | 68.33 | 55.88 | **$6.27\times10^{-6}$** | 423 | 80.58 | 65.81 | **0.0018** | **$2.99\times10^{-17}$** |
| Male Professors | 2093 | 78.39 | 61.69 | | 1060 | 93.45 | 72.32 | | **$1.31\times10^{-36}$** |

[a]Comparison of female vs. male faculty members affiliated with all 146 R1 universities.
[b]Comparison of female vs. male faculty members affiliated with the top 50 R1 universities.
[c]Comparison of faculty members affiliated with the top 50 R1 universities vs. faculty members affiliated with the other 96 R1 universities.
All *P*-values correspond to the Mann–Whitney *U* test. *P*-values lower than 0.05 are shown in bold face.

### Table S7: Average number of authors per publication

| | All 146 R1 universities | | | *P*-value[a] | Top 50 universities | | | *P*-value[b] | *P*-value[c] |
|---|---|---|---|---|---|---|---|---|---|
| **Gender and rank** | ***n*** | **Average** | **Median** | | ***n*** | **Average** | **Median** | | |
| Female Assistant Prof. | 628 | 8.52 | 6.91 | 0.1704 | 250 | 9.13 | 7.35 | 0.2559 | **0.0295** |
| Male Assistant Prof. | 710 | 9.40 | 7.23 | | 296 | 9.61 | 7.62 | | **0.0031** |
| Female Associate Prof. | 575 | 7.60 | 5.91 | 0.9829 | 178 | 8.92 | 6.15 | 0.3322 | **0.0084** |
| Male Associate Prof. | 918 | 7.74 | 6.03 | | 307 | 8.80 | 6.70 | | **$1.22\times10^{-7}$** |
| Female Professors | 901 | 7.61 | 5.64 | 0.1086 | 423 | 8.25 | 5.75 | 0.1579 | **0.0382** |
| Male Professors | 2093 | 7.06 | 5.47 | | 1060 | 7.45 | 5.60 | | **0.0087** |

[a]Comparison of female vs. male faculty members affiliated with all 146 R1 universities.
[b]Comparison of female vs. male faculty members affiliated with the top 50 R1 universities.
[c]Comparison of faculty members affiliated with the top 50 R1 universities vs. faculty members affiliated with the other 96 R1 universities.
All *P*-values correspond to the Mann–Whitney *U* test. *P*-values lower than 0.05 are shown in bold face.

### Table S8: Percent of publications as first author

| | All 146 R1 universities | | | *P*-value[a] | Top 50 universities | | | *P*-value[b] | *P*-value[c] |
|---|---|---|---|---|---|---|---|---|---|
| **Gender and rank** | ***n*** | **Average** | **Median** | | ***n*** | **Average** | **Median** | | |
| Female Assistant Prof. | 628 | 41.07 | 37.50 | **$6.73\times10^{-6}$** | 250 | 40.07 | 35.00 | **0.0044** | **0.0049** |
| Male Assistant Prof. | 710 | 35.76 | 33.33 | | 296 | 33.72 | 31.03 | | **0.0014** |
| Female Associate Prof. | 575 | 27.92 | 25.71 | 0.0917 | 178 | 24.11 | 22.73 | 0.3732 | **$1.94\times10^{-5}$** |
| Male Associate Prof. | 918 | 27.10 | 24.36 | | 307 | 23.16 | 20.99 | | **$5.90\times10^{-8}$** |
| Female Professors | 901 | 21.62 | 19.51 | 0.2031 | 423 | 20.21 | 18.39 | **0.0455** | **0.0002** |
| Male Professors | 2093 | 21.40 | 19.05 | | 1060 | 19.18 | 16.94 | | **$2.79\times10^{-16}$** |

[a]Comparison of female vs. male faculty members affiliated with all 146 R1 universities.
[b]Comparison of female vs. male faculty members affiliated with the top 50 R1 universities.
[c]Comparison of faculty members affiliated with the top 50 R1 universities vs. faculty members affiliated with the other 96 R1 universities.
All *P*-values correspond to the Mann–Whitney *U* test. *P*-values lower than 0.05 are shown in bold face.

### Table S9: Percent of publications as last author

| | All 146 R1 universities | | | *P*-value[a] | Top 50 universities | | | *P*-value[b] | *P*-value[c] |
|---|---|---|---|---|---|---|---|---|---|
| **Gender and rank** | ***n*** | **Average** | **Median** | | ***n*** | **Average** | **Median** | | |
| Female Assistant Prof. | 628 | 16.25 | 12.50 | 0.4229 | 250 | 17.74 | 15.38 | 0.1701 | **0.0239** |
| Male Assistant Prof. | 710 | 15.30 | 11.76 | | 296 | 16.57 | 11.76 | | 0.4356 |
| Female Associate Prof. | 575 | 34.88 | 35.00 | **0.0193** | 178 | 38.12 | 39.29 | **0.0026** | **0.0010** |
| Male Associate Prof. | 918 | 33.05 | 32.82 | | 307 | 33.79 | 33.82 | | 0.1426 |
| Female Professors | 901 | 47.29 | 47.93 | 0.9635 | 423 | 50.34 | 51.00 | 0.8261 | **$2.84\times10^{-7}$** |
| Male Professors | 2093 | 47.53 | 47.73 | | 1060 | 50.18 | 50.82 | | **$4.88\times10^{-16}$** |

[a]Comparison of female vs. male faculty members affiliated with all 146 R1 universities.
[b]Comparison of female vs. male faculty members affiliated with the top 50 R1 universities.
[c]Comparison of faculty members affiliated with the top 50 R1 universities vs. faculty members affiliated with the other 96 R1 universities.
All *P*-values correspond to the Mann–Whitney *U* test. *P*-values lower than 0.05 are shown in bold face.

### Table S10: Percent of publications as single author

| | All 146 R1 universities | | | *P*-value[a] | Top 50 universities | | | *P*-value[b] | *P*-value[c] |
|---|---|---|---|---|---|---|---|---|---|
| **Gender and rank** | ***n*** | **Average** | **Median** | | ***n*** | **Average** | **Median** | | |
| Female Assistant Prof. | 628 | 2.52 | 0.00 | 0.3848 | 250 | 2.33 | 0.00 | 0.5925 | 0.5385 |
| Male Assistant Prof. | 710 | 2.36 | 0.00 | | 296 | 2.70 | 0.00 | | 0.4647 |
| Female Associate Prof. | 575 | 3.74 | 1.47 | 0.0902 | 178 | 3.12 | 1.72 | 0.9154 | 0.7249 |
| Male Associate Prof. | 918 | 4.32 | 1.80 | | 307 | 3.21 | 1.49 | | **0.0293** |
| Female Professors | 901 | 5.13 | 3.23 | **0.0007** | 423 | 5.56 | 3.70 | 0.2789 | **0.0013** |
| Male Professors | 2093 | 5.97 | 3.70 | | 1060 | 5.97 | 3.85 | | 0.0609 |

[a]Comparison of female vs. male faculty members affiliated with all 146 R1 universities.
[b]Comparison of female vs. male faculty members affiliated with the top 50 R1 universities.
[c]Comparison of faculty members affiliated with the top 50 R1 universities vs. faculty members affiliated with the other 96 R1 universities.
All *P*-values correspond to the Mann–Whitney *U* test. *P*-values lower than 0.05 are shown in bold face.

### Table S11: Average impact factor of publications

| | All 146 R1 universities | | | *P*-value[a] | Top 50 universities | | | *P*-value[b] | *P*-value[c] |
|---|---|---|---|---|---|---|---|---|---|
| **Gender and rank** | ***n*** | **Average** | **Median** | | ***n*** | **Average** | **Median** | | |
| Female Assistant Prof. | 628 | 7.43 | 5.75 | **$2.39\times10^{-7}$** | 250 | 9.00 | 7.27 | **$1.31\times10^{-5}$** | **$1.01\times10^{-15}$** |
| Male Assistant Prof. | 710 | 8.69 | 7.20 | | 296 | 11.02 | 9.34 | | **$1.09\times10^{-23}$** |
| Female Associate Prof. | 575 | 6.22 | 5.38 | **0.0007** | 178 | 7.74 | 7.01 | **0.0064** | **$2.54\times10^{-15}$** |
| Male Associate Prof. | 918 | 6.96 | 6.12 | | 307 | 8.94 | 8.05 | | **$5.87\times10^{-26}$** |
| Female Professors | 901 | 7.06 | 6.07 | **0.0152** | 423 | 8.61 | 7.61 | **0.2205** | **$2.78\times10^{-32}$** |
| Male Professors | 2093 | 7.38 | 6.38 | | 1060 | 8.89 | 7.85 | | **$7.33\times10^{-74}$** |

[a]Comparison of female vs. male faculty members affiliated with all 146 R1 universities.
[b]Comparison of female vs. male faculty members affiliated with the top 50 R1 universities.
[c]Comparison of faculty members affiliated with the top 50 R1 universities vs. faculty members affiliated with the other 96 R1 universities.
All *P*-values correspond to the Mann–Whitney *U* test. *P*-values lower than 0.05 are shown in bold face.
Only publications published in journals indexed in the 2023 *Journal Citation Reports* were included I the analysis.

### SUPPLEMENTARY FIGURES

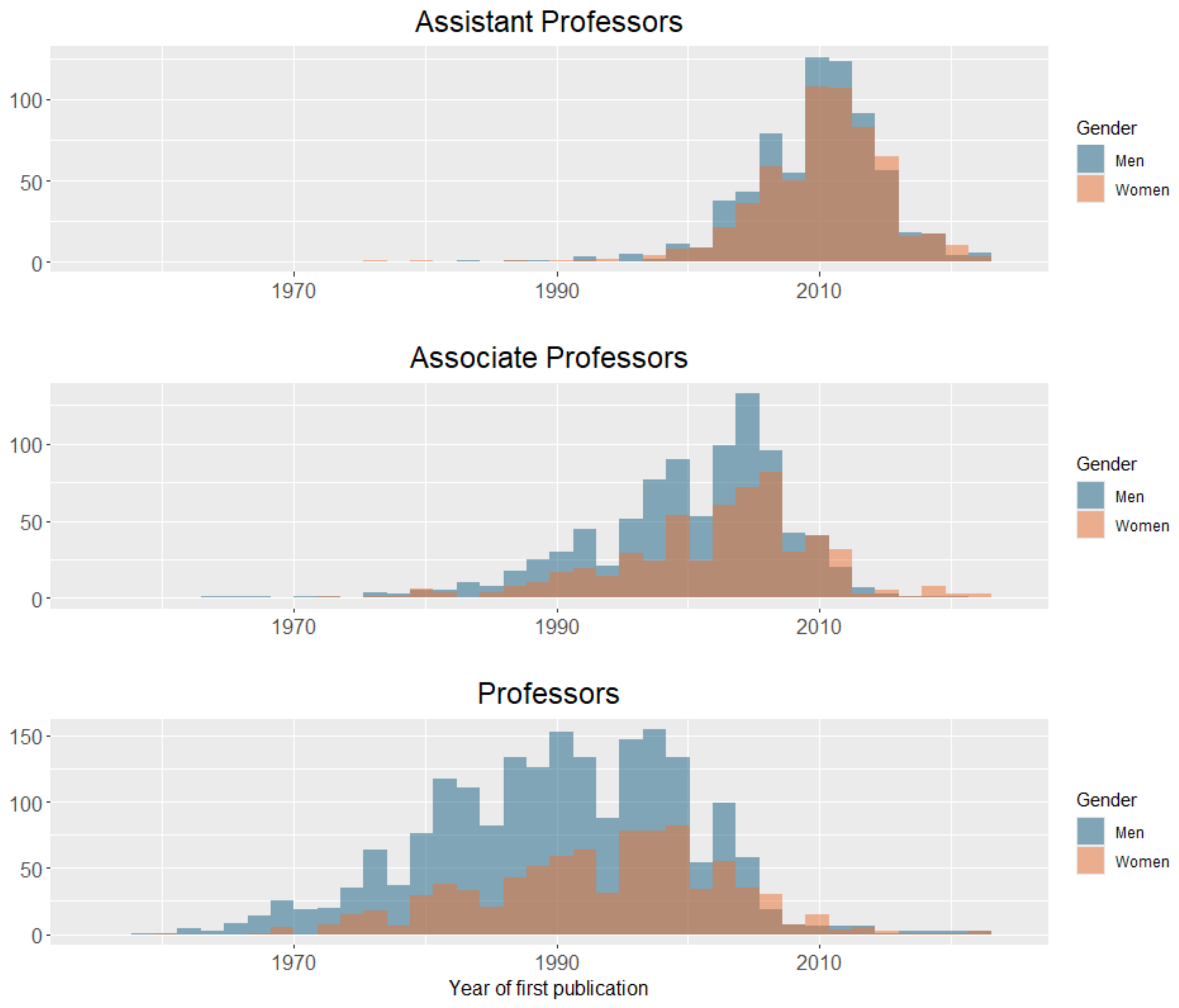


**Figure S1: Year of first publication of female and male faculty members of different ranks.**

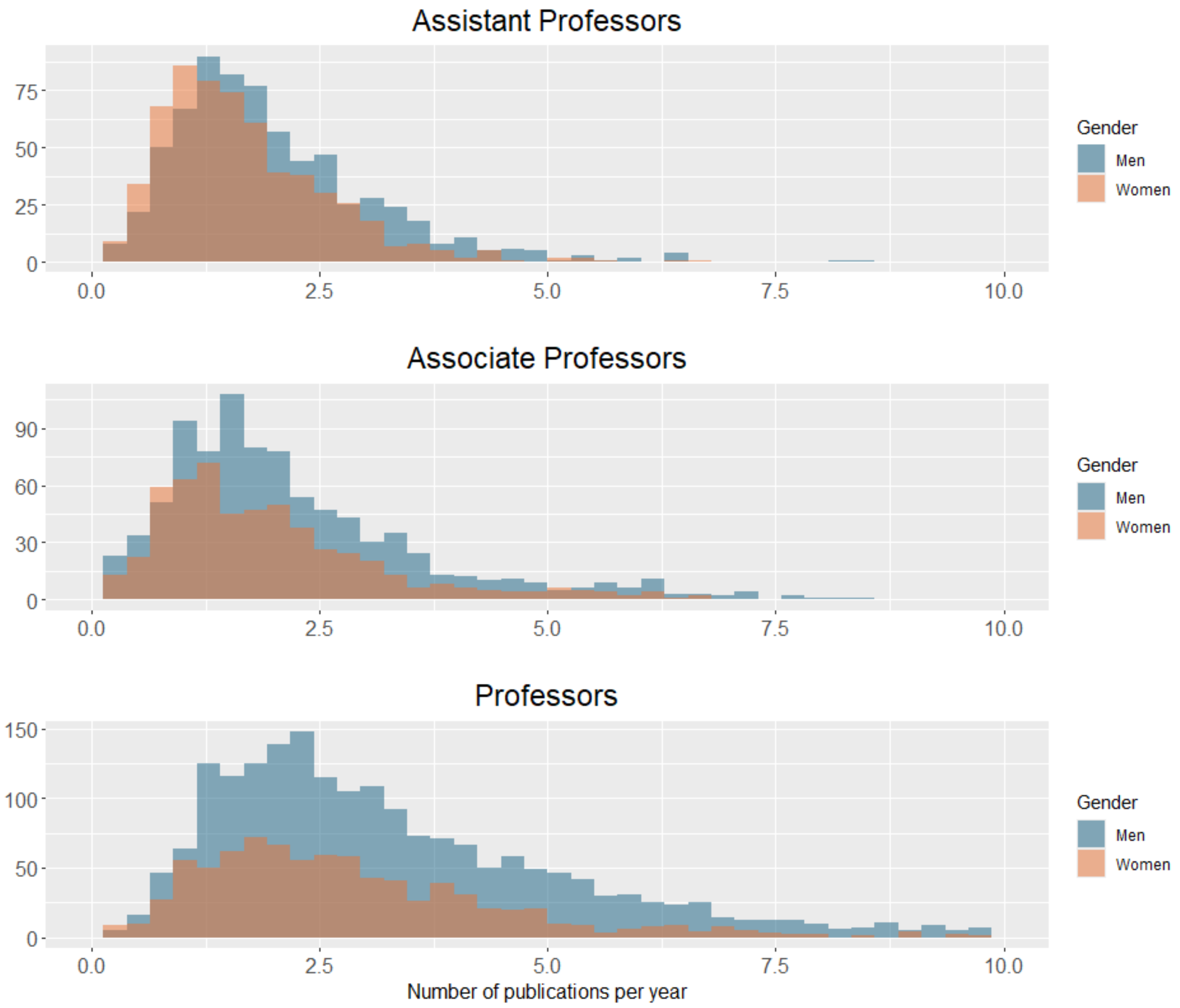


**Figure S2: Number of publications per year of female and male faculty members of different ranks.**

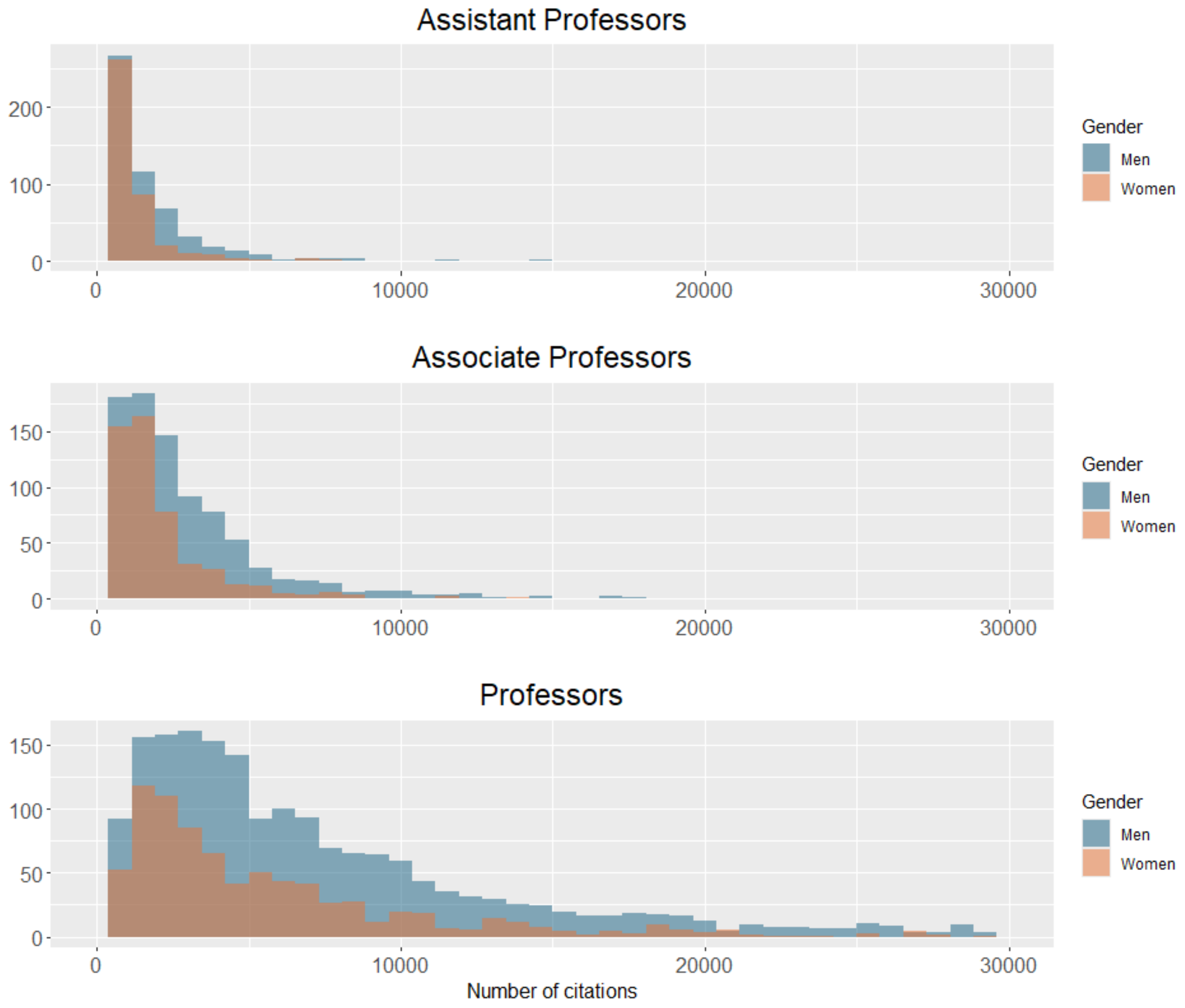


**Figure S3: Total number of citations of female and male faculty members of different ranks.**

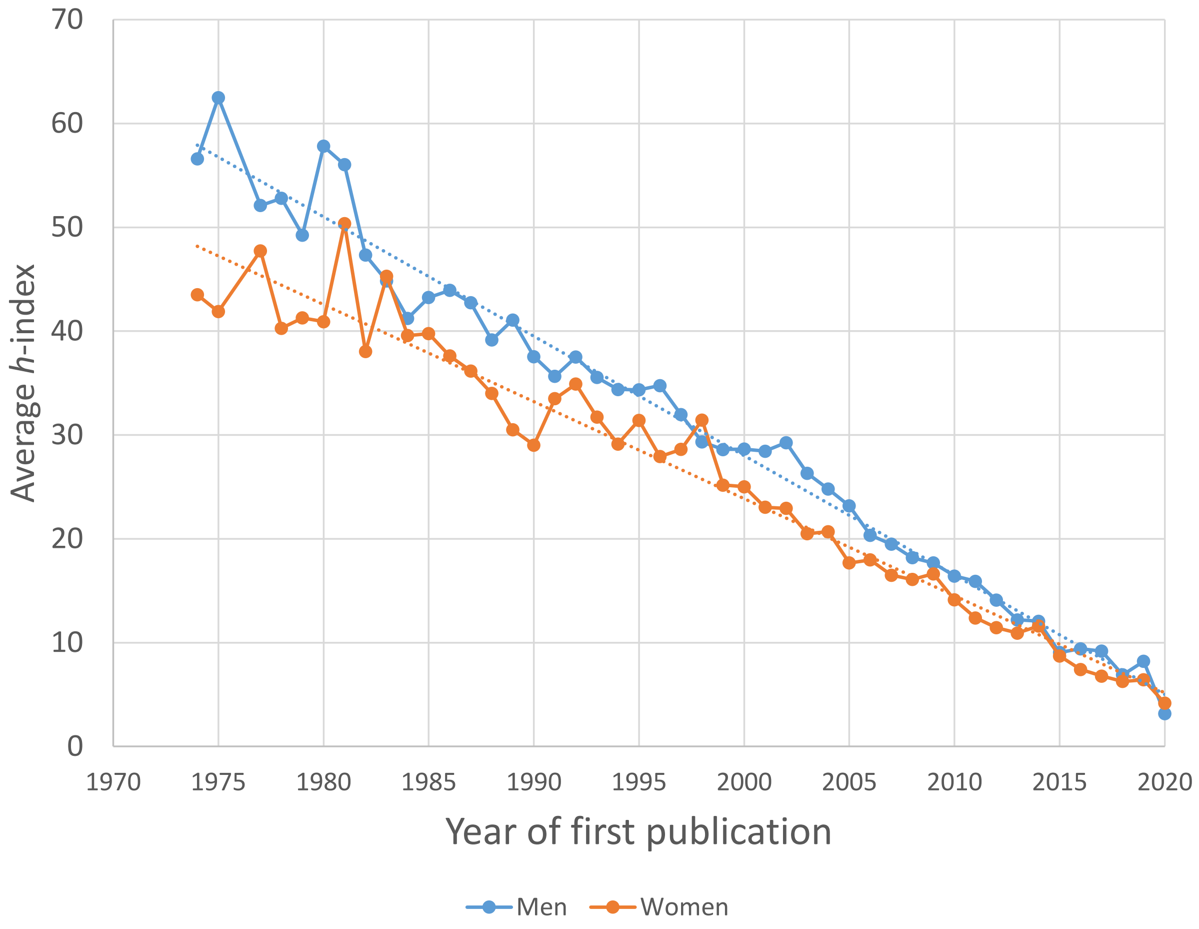


**Figure S4: Average *h*-index of women and men in each cohort.** Each cohort is composed of all faculty members that started publishing in a given year. Only cohorts with at least 6 women and 6 men are represented. Dotted lines represent regression lines.

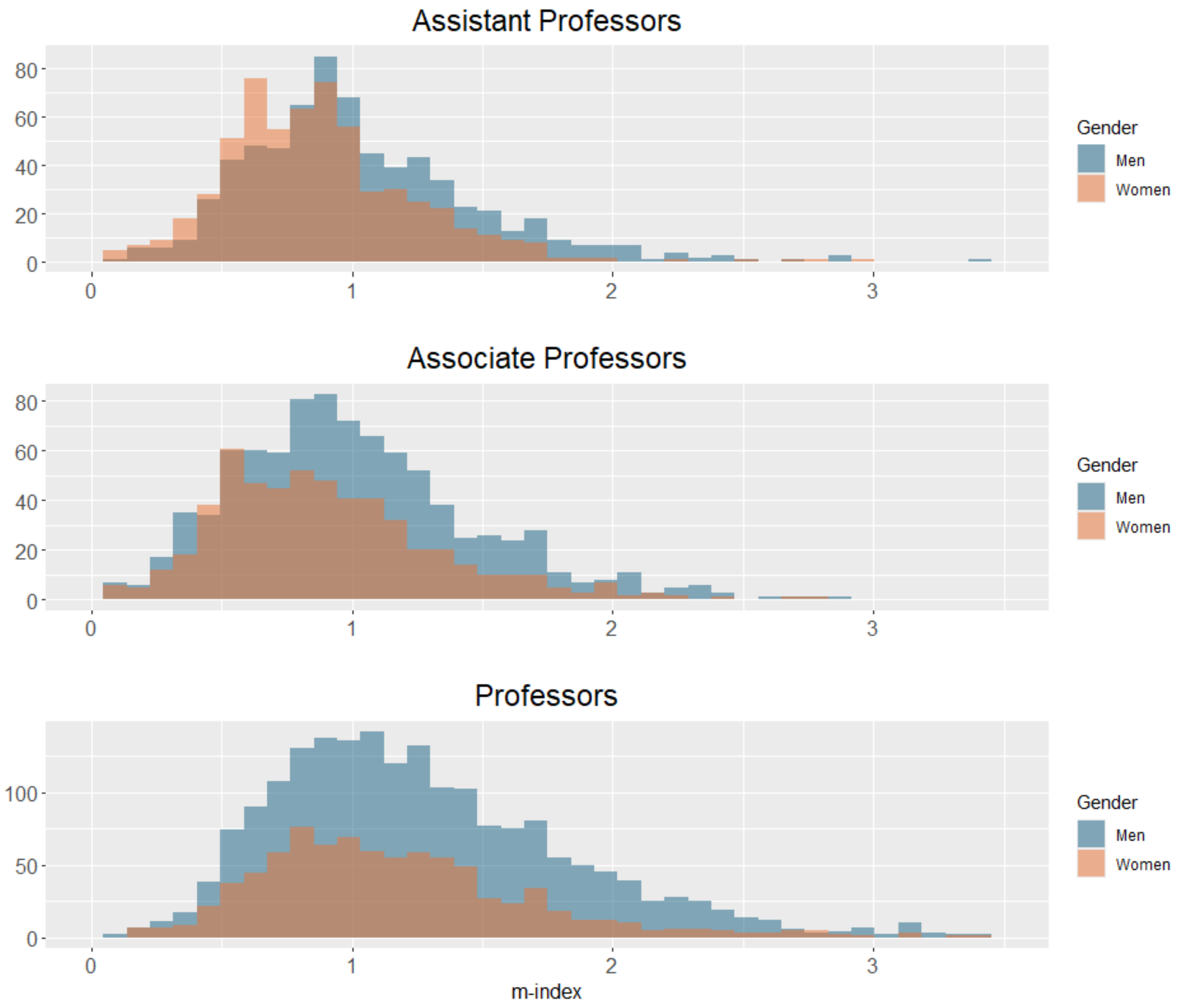


**Figure S5: *m*-index of female and male faculty members of different ranks.**

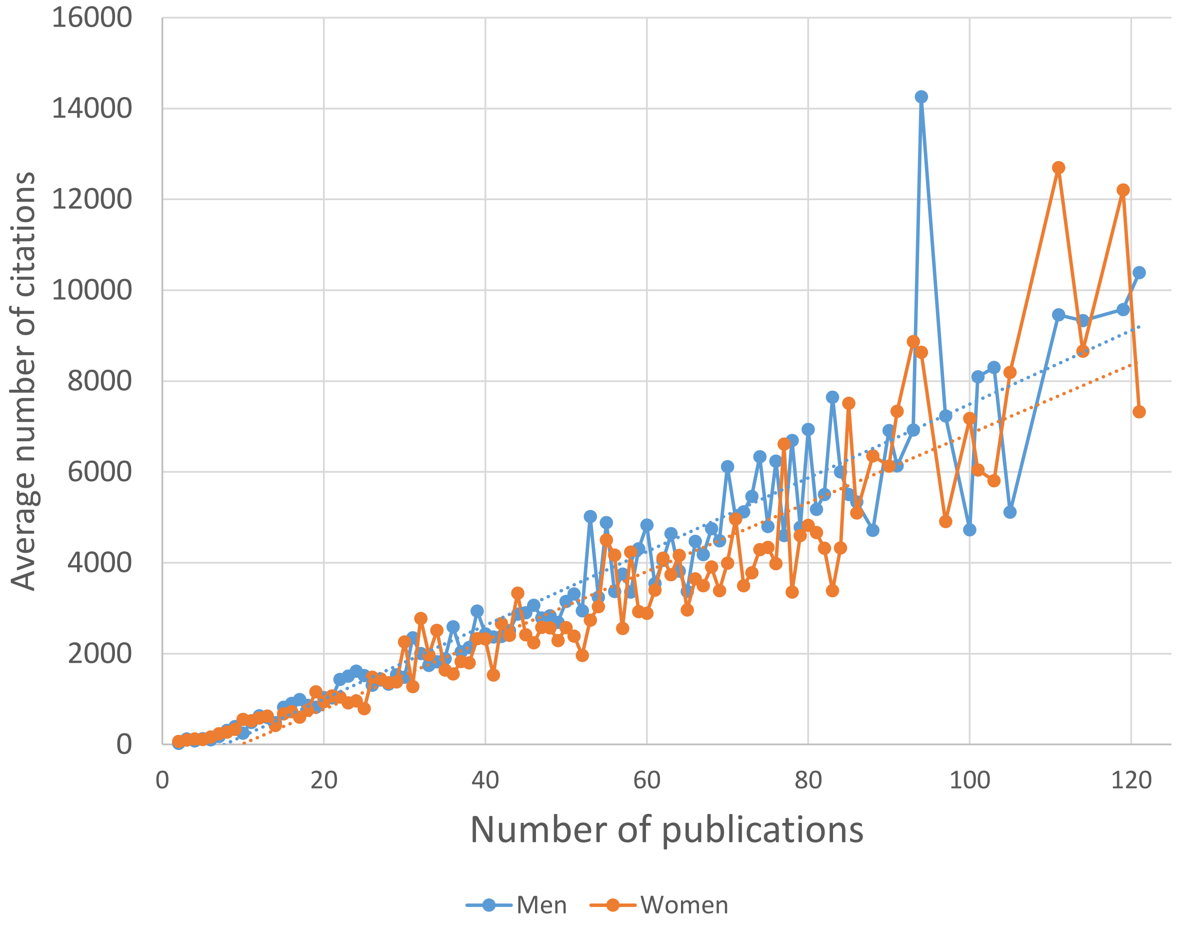


**Figure S6: Average number of total citations of women and men classified according to their number of publications.** Each group is composed of all faculty members that authored a given number of publications. Only groups with at least 6 women and 6 men are represented. Dotted lines represent regression lines.

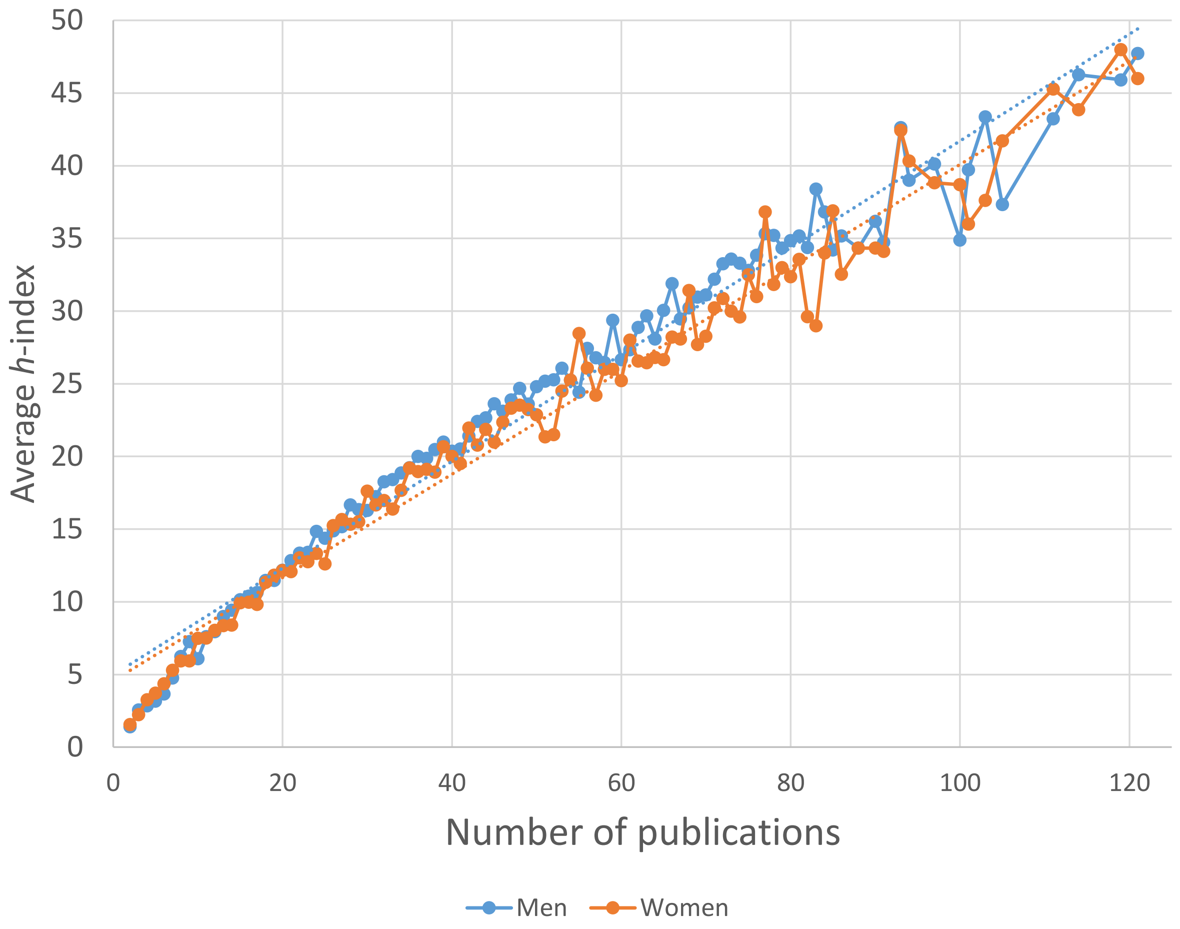


**Figure S7: Average *h*-index of women and men classified according to their number of publications.** Each group is composed of all faculty members that authored a given number of publications. Only groups with at least 6 women and 6 men are represented. Dotted lines represent regression lines.

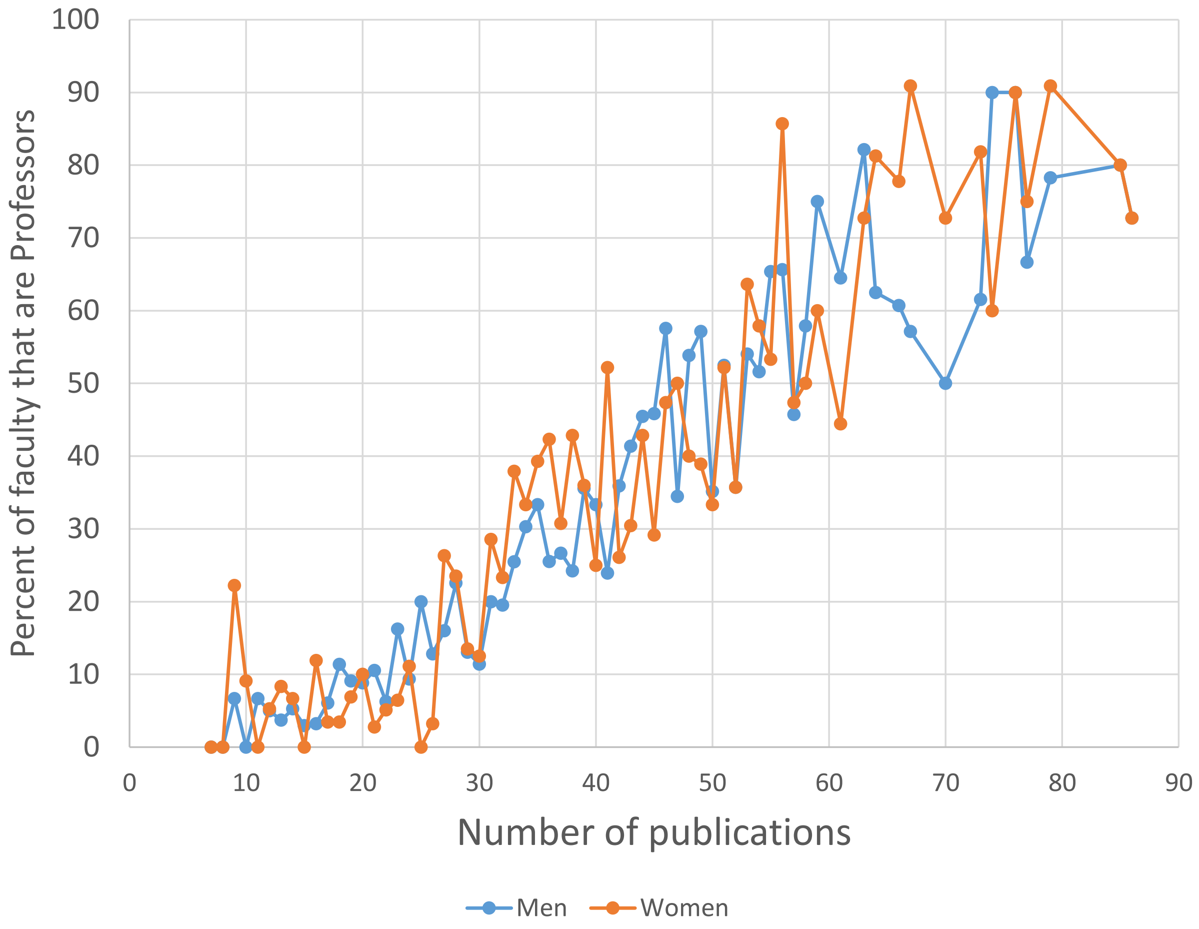


**Figure S8: Percentage of tenured and tenure-eligible faculty members that have attained the rank of Professor among women and men classified according to their number of publications.** Each group is composed of all faculty members that authored a given number of publications. Only groups with at least 10 women and 10 men are represented.

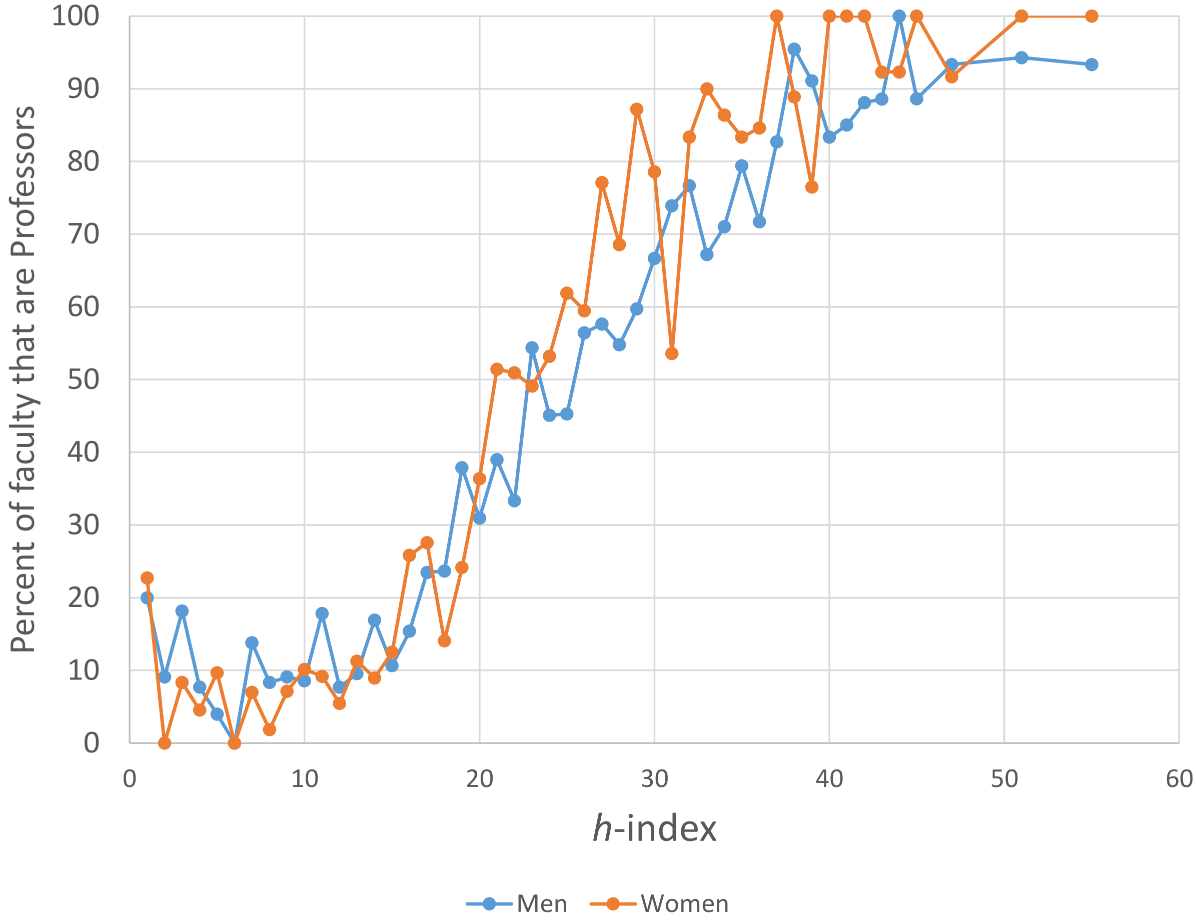


**Figure S9: Percentage of tenured and tenure-eligible faculty members that have attained the rank of Professor among women and men classified according to their *h*-index.** Each group is composed of all faculty members that shared the same *h*-index. Only groups with at least 10 women and 10 men are represented.
